# A *Sox2* Enhancer Cluster Regulates Region-Specific Neural Fates from Mouse Embryonic Stem Cells

**DOI:** 10.1101/2023.11.09.566464

**Authors:** Ian C Tobias, Sakthi D Moorthy, Virlana M Shchuka, Lida Langroudi, Mariia Cherednychenko, Zoe E Gillespie, Andrew G Duncan, Ruxiao Tian, Natalia A Gajewska, Raphaël B Di Roberto, Jennifer A Mitchell

## Abstract

Embryonic development depends on spatially and temporally orchestrated gene regulatory networks. Expressed in neural stem and progenitor cells (NSPCs), the transcription factor sex-determining region Y box 2 (Sox2) is critical for embryogenesis and stem cell maintenance in neural development. Whereas *Sox2* is regulated by a distal cluster of enhancers in embryonic stem cells (ESCs), enhancers closer to the gene have been implicated in *Sox2* transcriptional regulation in the neural lineage. Using functional genomics data, and deletion analysis we show that a downstream enhancer cluster regulates *Sox2* transcription in NSPCs derived from mouse ESCs. By generating allelic mutants using CRISPR-Cas9 mediated deletions, we show that this proximal enhancer cluster, termed *Sox2* regulatory regions 2-18 (SRR2-18), is a *cis* regulator of *Sox2* transcription during neural differentiation. Transcriptome analyses demonstrate that loss of even one copy of SRR2-18 disrupts the region-specific identity of NSPCs. Biallelic deletion of this *Sox2* neural enhancer cluster causes reduced SOX2 protein, less frequent interaction with transcriptional machinery, and leads to perturbed chromatin accessibility genome-wide further affecting the expression of neurodevelopmental and anterior-posterior regionalization genes. Furthermore, homozygous NSPC deletants exhibit self-renewal defects and impaired differentiation into cell types found in the brain. Altogether, our data define a *cis*-regulatory enhancer cluster controlling *Sox2* transcription in NSPCs and highlight the sensitivity of neural differentiation processes to decreased *Sox2* transcription, which influences their differentiation into posterior neural fates, specifically the caudal neural tube.

## Introduction

Specialized cell types within the nervous system are generated during embryonic development from populations of self-renewing stem and progenitor cells that emerge from defined neurogenic zones (Johe et al. 1996; Molofsky et al. 2003). These neural stem and progenitor cells (NSPCs) migrate and undergo differentiation to create interconnected cellular systems consisting of neuronal and glial cell types, which include astrocytes and oligodendrocytes (Götz and Barde 2005; Shen et al. 2006). More than 80 different neuronal and glial cell types corresponding to developmental stage, regional specificity, and cellular function have been identified by single cell gene expression patterns in the mouse postnatal brain and spinal cord (Rosenberg et al. 2018; Zeisel et al. 2018). To execute the complex functions of the vertebrate nervous system, regionally distinct gene expression profiles instruct NSPC populations to organize specialized cell types in the brain and spinal cord along the rostral-caudal and dorsal-ventral axes (Mathis and François Nicolas 2000; Hagey et al. 2016).

The proliferative activity, developmental potency, and region-specific identity of NSPCs are controlled by genetic programs defined by differential regulatory element usage (Ensini et al. 1998; Ziller et al. 2015). Cell fate decisions involve extensive transcriptional changes that are governed by gene regulatory networks — functional DNA sequences (such as enhancers) responsive to transcription factors (TFs), chromatin topology, and/or epigenetic modifiers (Brandenberger et al. 2004; Loh et al. 2006; Bailey et al. 2006). Enhancers are orientation-independent regulatory elements that influence when and where genes become transcriptionally active (Banerji et al. 1981). In models that explore cell differentiation processes, coordinated changes in TF binding occur at enhancer regions, which directly impact target gene expression and restrict cellular developmental outcomes (Wamstad et al. 2012; Kieffer-Kwon et al. 2013; Huang et al. 2016). Enhancers can be positioned upstream or downstream and at variable genomic distances from the genes they regulate (Tuan et al. 1989; Sanyal et al. 2012). Depending on their genomic location, enhancers have been shown regulate gene transcription through the formation of specific chromatin-chromatin interactions (Carter et al. 2002; Tolhuis et al. 2002; Zhang et al. 2013). Distal enhancer elements can be targeted to promoters by architectural elements, either through the combined action of the zinc finger protein CCCTC-binding factor (CTCF) and the ring-shaped Cohesin Complex, or by CTCF-independent mechanisms (Tang et al. 2015; Weintraub et al. 2017; Taylor et al. 2022).

Enhancer sequence-containing regions exhibit a variety of functionally-correlated chromatin states, which are often inferred from DNA methylation and post-translational histone modifications (ENCODE Project Consortium 2012). Additionally, enhancers appear to be more dynamic than promoter elements across various cell types, tissues, and taxonomic groups (Heintzman et al. 2009; Villar et al. 2015; Roadmap Epigenomics Consortium et al. 2015). By leveraging our knowledge of epigenetically-informed chromatin states (e.g., Histone H3 lysine 27 acetylation [H3K27Ac] in the context of enhancers), TF occupancy, chromatin accessibility, and co-activator recruitment (e.g., Mediator complex, EP300) (Kagey et al. 2010; Creyghton et al. 2010; Rada-Iglesias et al. 2011), we can make predictions about enhancer activity. However, a single gene can be regulated by multiple enhancers active in different cellular contexts or by redundant enhancers with some degree of overlapping spatiotemporal function (Hay et al. 2016; Moorthy et al. 2017; Osterwalder et al. 2018; Taylor et al. 2022). As a result, identifying the locations of enhancers and determining their phenotypic role(s) is an ongoing challenge (reviewed in (Tobias et al. 2021)).

Sex-determining region Y box 2 (*Sox2*) encodes a TF that is initially expressed in the inner cell mass and epiblast of preimplantation embryos (Gubbay et al. 1990; Avilion et al. 2003). *Sox2* expression is downregulated upon pluripotent cell commitment towards endodermal or mesodermal lineages; however, its transcription is maintained within the neuroectodermal lineage, where SOX2 inhibits mesendoderm fate determinants (Ivanova et al. 2006; Kopp et al. 2008; Thomson et al. 2011). *Sox2* represents one of the SOXB1 family transcription factors (*Sox1-3*) which have partially overlapping DNA-binding activity in the neural ectoderm and the anterior gut endoderm (Taranova et al. 2006; Que et al. 2007; Cavallaro et al. 2008). SOX2 functions collaboratively with other TFs, including *Pou5f1* and *Pax6*, to ensure proper regulation of transcriptional processes that are essential for maintaining tissue-specific identities (Masui et al. 2007; Chen et al. 2008; Lodato et al. 2013; Quevedo et al. 2019).

The deletion of *Sox2* in the E10.5-12.5 forebrain produces viable mice with mild neurodevelopmental impairments at birth (Miyagi et al. 2008). Conversely, *Sox2* is essential for neurogenesis in the adult mouse hippocampal niche, which exhibits a Sonic hedgehog (Shh) signaling-dependent depletion of NSPCs in the absence of *Sox2* (Favaro et al. 2009). NSPCs display regional and temporal heterogeneity *in vivo*, with both quiescent and amplifying progenitor cell populations that retain self-renewal and differentiation capabilities when cultured *in vitro* (Reynolds and Weiss 1992; Garcia et al. 2004; Suh et al. 2007). Diffusible factors can induce NSPCs to shift positional neural cell identities along the rostral-caudal or ventral-dorsal axes; however, the primary allocation of embryonic cells to the anterior neuroepithelium which derives the brain and cervical spinal cord progenitors versus ‘posteriorized’ neural progenitors that drive elongation of the neural tube is thought to occur in the vertebrate epiblast prior to neural induction (Bertacchi et al. 2013; Hallmann et al. 2016; Metzis et al. 2018). These forms of regionalized gene regulation in neural fated progenitor populations are essential to prevent the specification of inappropriate or mixed cell identities.

*Sox2* and its genic flanking sequences fall within a synteny block on chromosome 3 (q26.33) in placental mammals (Zhang et al. 2022). Multiple regulatory elements within this conserved sequence block show regionally biased activities and are posited to coordinate *Sox2* transcription in diverse neurogenic contexts where SOX2 is required (Uchikawa et al. 2003; Uchikawa and Kondoh 2016). In mice, two enhancers located proximally to the *Sox2* transcriptional start site (TSS), known as *Sox2* regulatory regions (SRR) 1 and 2, have been shown to independently drive reporter gene expression in both ESCs and NSPCs (Zappone et al. 2000; Tomioka et al. 2002; Miyagi et al. 2004; Catena et al. 2004). Biallelic deletion of SRR1 in mouse embryos has been shown to induce a transient *Sox2* expression deficit in the anterior neural plate (Iwafuchi-Doi et al. 2011). *Sox2* haplodeficient mice with a monoallelic deletion of SRR1 on the intact *Sox2* allele display cerebral malformations and a decrease in Nestin (*Nes*)-positive hippocampal NSPCs (Ferri et al. 2004). Conversely, SRR2 is thought to recruit transcriptional repressors and decrease *Sox2* expression during *in vitro* differentiation of embryonic and neural stem cells (Li et al. 2012; Marqués-Torrejón et al. 2013). Another *Sox2* regulatory element homologous to the N1 enhancer in chickens is first activated in the caudal lateral epiblast of mouse embryos and later controls the fate of *Sox2*-expressing neuromesodermal progenitor (NMP) cells (Takemoto et al. 2006; Takemoto et al. 2011). However, there still exists a significant gap in understanding between the identification of *Sox2* regulatory elements and how the interplay between distinct *cis*-regulatory elements allows for the fine-tuning of transcription required for specific developmental fates.

The ESC-to-NSPC differentiation model provides a genetically tractable system for studying how epigenetic remodeling and chromatin organization influence neural gene regulatory networks (Phillips-Cremins et al. 2013). We have shown that the *Sox2* control region (SCR), a distal cluster of enhancers located at position +104-111 kb downstream of the *Sox2* TSS, regulates *Sox2* expression in mouse ESCs (Chen et al. 2012; Zhou et al. 2014). As pluripotent cells differentiate, TF binding is redistributed genome-wide and the chromatin state is altered at the SCR, including a loss of H3K27ac enrichment and long-range interactions with the *Sox2* promoter (Beagan et al. 2016; Thompson et al. 2022; Chakraborty et al. 2023). Leveraging the context-dependent activity of this model enhancer cluster, the *Sox2* locus has been extensively used in various studies that investigated long-range gene regulation in pluripotent stem cells (Brosh et al. 2023; Platania et al. 2023; Chakraborty et al. 2023). However, a more comprehensive understanding of how regulatory elements contribute to *Sox2* gene regulation in neural lineage commitment is still required. To test *Sox2* regulatory mechanisms in neural cells, we investigated the contribution of candidate neural lineage *Sox2* enhancers using mouse ESC-derived NSPCs and CRISPR-Cas9 mediated genome engineering.

## Results

### *Sox2* proximal regions are responsible for enhancement of *Sox2* transcription during differentiation to neural stem/progenitor cells

SRR1 and SRR2 exhibit enhancer activity in the mouse epiblast and in ESCs, suggesting their potential role in *Sox2* gene regulation early in development (Iwafuchi-Doi et al. 2011). Neither of these enhancers produce an obvious neurodevelopmental phenotype when both allelic copies are deleted in non-sensitized (*Sox2*^+/+^) genetically engineered mouse models (Ferri et al. 2004; Moncho-Amor et al. 2021). This raises the possibility that additional *Sox2* enhancer elements, including those already known to be conserved in amniotes that may be under selective pressure, could have partially overlapping or compensatory spatiotemporal activities that are not captured by conventional ectopic enhancer-reporter assays (Uchikawa et al. 2003). We reanalyzed chromatin immunoprecipitation with sequencing (ChIP-seq) data from the ENCODE consortium to examine the coactivator-associated modification H3K27ac in mouse ESCs and E14.5 forebrain tissue as an epigenetically informed approach to identify enhancer candidates with similar activity patterns during early brain development (**Fig. 1A**; ENCODE dataset accession numbers provided in **Table S1**) (ENCODE Project Consortium 2012). To refine our search to regions contributing to RNA polymerase II (RNAPII)-mediated transcription, we also reanalyzed Mediator subunit MED1 ChIP-seq data from *in vitro* generated NSPCs (SRA and ENA dataset accession numbers provided in **Table S2**) (Quevedo et al. 2019).

**Fig 1.**
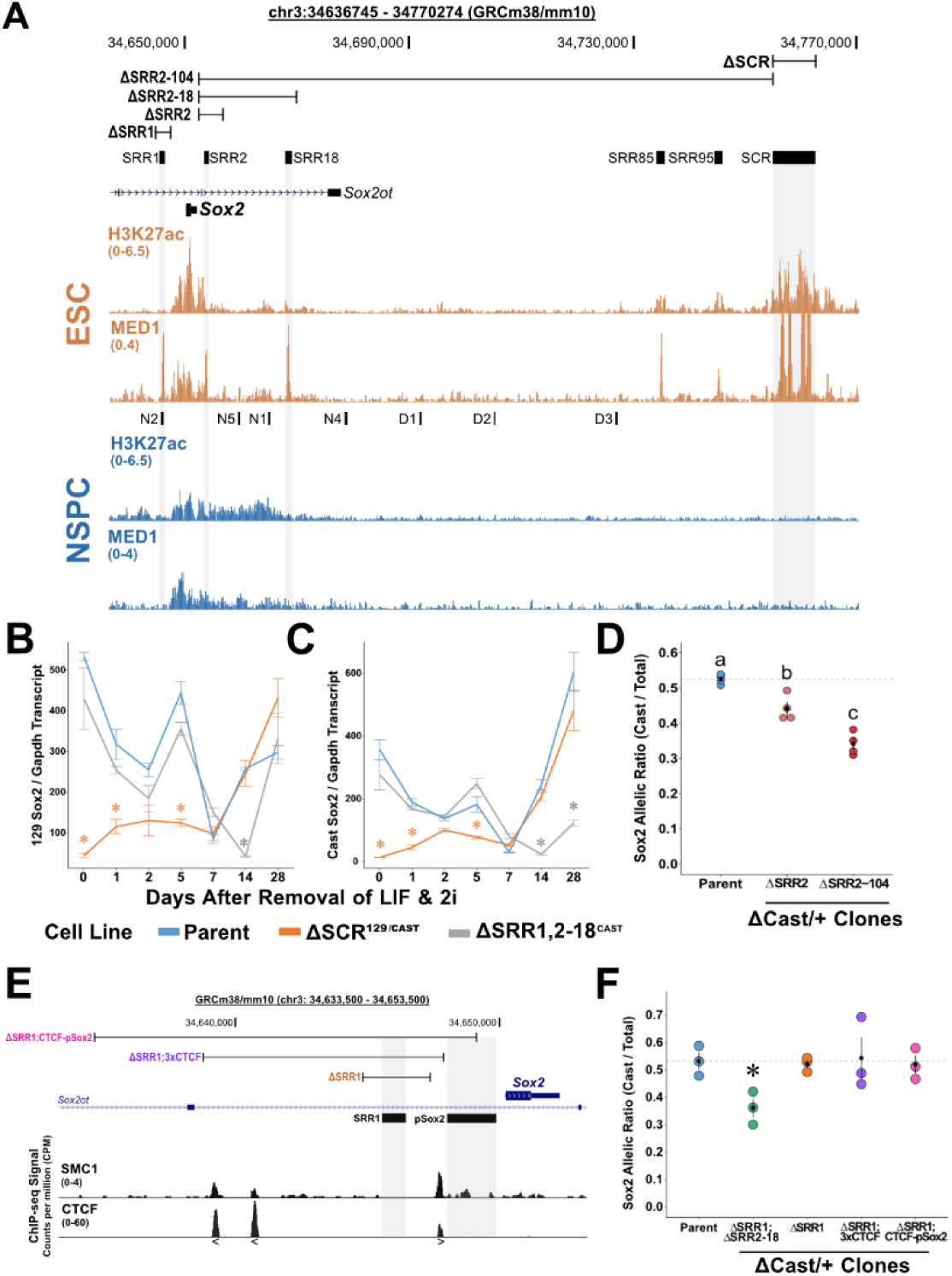
*Sox2* enhancer switch during mouse ESC neural differentiation. **(A)** H3K27ac and MED1 ChIP-seq tracks in mouse embryonic stem cells (mESCs; orange) and neural stem/progenitor cells (NSPCs; blue) visualized by UCSC Genome Browser (mm10). Black boxes aligned with shaded bars mark *Sox2* regulatory regions (SRRs) active in mESCs. Ranges show the margins of CRISPR-Cas9-mediated deletions of different SRRs or the *Sox2* control region (SCR). **(B-C)** *Sox2* expression in ESCs undergoing neural induction by allele-specific RT-qPCR comparing wildtype lines to lines harboring monoallelic (SRR1,2-18) or biallelic (SCR) deletions of enhancers with the indicated genotypes. Significantly different groups from the day 0 timepoint are indicated (∗) *P* < 0.05. Error bars represent SEM. n ≥ 3. **(D)** *Sox2* expression in stably derived NSPCs by allele-specific RT-qPCR comparing wildtype lines to lines harboring monoallelic deletions of enhancers with the indicated genotypes. Expression levels for each allele are shown relative to the total transcript levels. Groups determined to be significantly different (*P* < 0.05) from one another are labeled with different letters; to indicate *P* > 0.05, groups would be labeled with the same letter. **(E)** SMC1 and CTCF ChIP-seq tracks visualized by the UCSC Genome Browser (mm10). Ranges show the margins of CRISPR-Cas9 mediated deletions of SRR1, clustered CTCF binding sites (3x CTCF) or the *Sox2* promoter (pSox2). Angled brackets indicate CTCF motif orientation. **(F)** *Sox2* expression from the Cast allele in ESC-derived NSPCs by allele-specific RT-qPCR comparing wildtype lines to lines harboring monoallelic deletions of enhancers with the indicated genotypes. Expression levels for each allele are shown relative to the total transcript level. Significant differences from the parent line values are denoted (∗) P < 0.05. Error bars represent SEM. n ≥ 3.

Examination of these ChIP-seq profiles showed that ESCs and NSPCs display distinct regions of H3K27ac enrichment surrounding the *Sox2* gene (**Fig. 1A**). The NSPC H3K27ac interval extends from the upstream flanking region, through SRR1, and into a 16.7 kb region downstream of the *Sox2* TSS containing SRR2 (we termed this downstream region SRR2-18) (**Fig. 1A**). MED1 chromatin recruitment signals were largely restricted to a 10 kb region immediately downstream of *Sox2* in cultured neural progenitors (**Fig. 1A**). We also observed H3K27ac enrichment in the SRR2-18 region in pooled E12.5 ganglionic eminence tissue from published forebrain epigenomic data (Rhodes et al. 2022) (**Supplemental Fig. S1A**), representing transient neural progenitor-rich niche regions in the ventral telencephalon that produce multiple neuronal subtypes in the cerebral cortex. These profiles of transcriptional co-activator activity suggested that *Sox2*-flanking regions, and SRR2-18 in particular, display enhancer associated features in multiple neurogenic niches containing telencephalic NSPCs.

To investigate gene expression dynamics during neural differentiation, we modeled neural induction and NSPC generation using mouse hybrid F1 (*M. musculus*/129 × *M. castaneus*) ESC cultures. To validate this system, we used RT-qPCR to measure the levels of *Sox2*, *Pou5f1*, *Pax6* and *Nes* RNA over the differentiation time course. We observed a gradual decrease in *Pou5f1* transcript levels as mouse ESCs transitioned from states of naïve pluripotency to the neural lineage in neural stem expansion (NSE) medium (*P* = 1.53 × 10^-2^, Wilcoxon Test; **Supplemental Fig. S1B**). Conversely, *Pax6* expression was significantly increased by day 5 of neural induction (*P* = 3.42 × 10^-5^, Welch’s T Test; **Supplemental Fig. S1C**), consistent with its role as a neural fate determinant (Suter et al. 2009). Expression of NSPC marker *Nes* (Lendahl et al. 1990) steadily increased with culturing in NSE medium (*P* = 1.21 × 10^-5^, Welch’s T Test; **Supplemental Fig. S1D**), whereas *Sox2* RNA levels peaked earlier at 3 days of neural induction (*P* = 2.47 × 10^-2^, Welch’s T Test; **Supplemental Fig. S1E**). Immunofluorescence imaging of monoclonal NSPC lines established from F1 ESCs confirmed the co-expression of both SOX2 and NES (**Supplemental Fig. S1F**).

We asked whether the combination of SRR1 and SRR2-18 play a role in coordinating *Sox2* expression during neural lineage commitment. Previous CRISPR-Cas9 mediated deletion analyses by our group in mouse F1 ESCs established two different enhancer-deleted cell lines: one lacking both allelic copies of the SCR (ΔSCR^129/Cast^), and the other harboring monoallelic compound deletions of candidate enhancers located within 18 kb of the *Sox2* TSS (ΔSRR1,2-18^Cast^) (Zhou et al. 2014). The latter deletion encompasses most of the neural H3K27ac domain flanking the *Sox2* gene. To monitor *Sox2* expression from each allele and across a range of *Sox2* transcription levels, we used allele-specific RT-qPCR to determine the proportion of the sum *Sox2* transcript quantity detected from the 129 or Cast allele, as we have done previously (Moorthy et al. 2017). In line with our previous finding that reduced *Sox2* dosage in ΔSCR^129/Cast^ ESCs delays but does not abolish pan-neural progenitor gene activation during undirected differentiation (Zhou et al. 2014), *Sox2*-deficient ΔSCR^129/Cast^ cells showed an increase in *Sox2* expression during neural differentiation (*P* = 3.73 × 10^-2^, Wilcoxon Test; **Fig. 1B**), and reached a level that was not significantly different from the unmodified control line at differentiation day 7 and in stable NSPC cultures (*P* =0.46, Wilcoxon Test; **Fig. 1B**). We did not observe any change in Cast *Sox2* transcript levels in ΔSRR1,2-18^Cast^ ESCs compared to the parent F1 ESCs and during the first two days of neural differentiation (*P* = 0.46 and *P* = 0.88, respectively, Wilcoxon Tests; **Fig. 1C**).

This is consistent with our previous finding that the *Sox2* proximal elements are dispensable for *Sox2* expression in pluripotent mouse ESCs (Zhou et al. 2014). Interestingly, ΔSRR1,2-18^Cast^ NSPCs showed decreased Cast-derived *Sox2* RNA compared to the unmodified, differentiation stage-matched controls (*P* = 2.86 × 10^-2^, Wilcoxon Test; **Fig. 1C**). These findings indicate that gene proximal elements within the SRR1 and/or SRR2-18 regions contribute to ESC-derived NSPC *Sox2* expression.

We next asked whether any additional enhancer elements within the downstream flanking region function in combination with SRR2 to activate *Sox2* transcription in our NSPC model system. To this end, we generated clonal populations of ESCs harboring a large deletion of the ∼100 kb region spanning SRR2 and SRR104 (ΔSRR2-104^Cast^), a mutant generated to remove all the possible downstream enhancers between SRR2 and the SCR (**Fig. 1A**). We also established NSPCs from mutant ESCs with a monoallelic deletion of SRR2 alone (ΔSRR2^Cast^) and assessed allele-specific *Sox2* transcript levels. This allowed us to distinguish between the contribution of SRR2, an element with reported activity in epiblast stem cell-derived neural progenitors (Iwafuchi-Doi et al. 2011), and other known or novel elements within the SRR2-104 region (e.g. N5, N1, N4) (**Fig 1A**). NSPC clones with either ΔSRR2^Cast^ or ΔSRR2-104^Cast^ genotypes showed a significant decrease in *Sox2* transcription from the targeted allele (*P* = 1.39 × 10^-2^ and *P* = 4.46 × 10^-4^, respectively, Tukey’s Test; **Fig. 1D**). Additionally, the proportion of Cast-derived *Sox2* transcript was further decreased in ΔSRR2-104^Cast^ cells compared to that in ΔSRR2^Cast^ NSPCs (*P* = 6.31 × 10^-3^, Tukey’s Test). Together, these data demonstrate that SRR2 and one or more regions downstream of SRR2 contribute to *Sox2* expression in ESC-derived NSPCs.

The architectural domain harboring *Sox2* in pluripotent and neural cells is delimited on the centromeric end by three CTCF-bound regions upstream of the *Sox2* promoter. We next focused on these upstream CTCF-bound sites that bookend SRR1, an enhancer in the anterior neural plate and in the dorsal telencephalon at mid-gestation (Zappone et al. 2000; Iwafuchi-Doi et al. 2011). To scrutinize the protein-binding elements with potential architectural and regulatory roles upstream of *Sox2*, we used ENCODE ChIP-seq data that profile CTCF binding in E14.5 mouse forebrain tissue (ENCODE Project Consortium 2012). We also reanalyzed publicly available ChIP-seq data for Cohesin complex subunit structural maintenance of chromosomes 1 (SMC1) retention in purified bulk NSPCs (Phillips-Cremins et al. 2013). Each of the CTCF-bound regions upstream of *Sox2* showed SMC1 accumulation in NSPCs (**Fig. 1E**), raising the possibility that local chromatin topology contributes to *Sox2* expression in neural progenitors.

To test whether SRR1 and the promoter proximal CTCF sites are required for *Sox2* expression in our ESC-derived NSPC model, we deleted these upstream regions encompassing SRR1 in F1 ESCs and differentiated genetically modified clones into NSPCs. As an external control, we confirmed the *cis*-linked decrease in *Sox2* allelic ratio in NSPCs established from ΔSRR1,2-18^Cast^ ESCs (*P* = 2.86 × 10^-2^, Welch’s T Test; **Fig. 1F**). Deletion of SRR1 on the Cast allele (ΔSRR1^Cast^) did not significantly change the allelic ratio of *Sox2* transcripts compared to unmodified control NSPCs (*P* = 0.77, Welch’s T Test; **Fig. 1F**). Moreover, we expanded the boundary of the SRR1-deleted region to include all three upstream CTCF binding sites in mouse neural cells (ΔSRR1-3xCTCF^Cast^). We did not observe any significant difference in *Sox2* allelic ratio in ΔSRR1-3xCTCF^Cast^ NSPCs (*P* = 0.90, Welch’s T Test; **Fig. 1F**), which is consistent with our recent finding that disruption of *Sox2* promoter proximal CTCF sites in the E9.5-10.5 mouse embryo does not affect SOX2 levels in the central nervous system (CNS) (Chakraborty et al. 2023). We have found that genes transactivated by regulatory architectures similar to the array of modular enhancers found within the *Sox2* TAD can have partially redundant regulatory activities; and may be individually dispensable for the maintenance of robust developmental gene expression (Moorthy et al. 2017). It also remained a possibility that the extended CpG island promoter region contributes to strong baseline levels of *Sox2* expression in neural progenitors. Yet after establishing NSPC clones with a larger deletion (-15.6 kb to -1.1 kb of TSS) that spanned all proximal CTCF binding sites, SRR1 and greater than half of the CpG island promoter region on the Cast allele (ΔSRR1 CTCF-pSox2^Cast^), the fraction of *Sox2* transcript from the Cast allele did not significantly differ from that of control NSPCs (*P* = 0.79, Welch’s T Test; **Fig. 1F**). In summary, our data suggest neither SRR1, the CTCF binding sites, nor the full-length CpG island promoter are required for *Sox2* transcription in this ESC model of neural lineage commitment.

### SRR2-18 is a *cis*-regulator of *Sox2* transcription and monoallelic deletion disrupts transcriptional regulatory programs in NSPCs

Since the H3K27ac- and MED1-enriched chromatin profiles in NSPCs were concentrated into a genomic window larger than a typical single active enhancer which are often predictive of spatially clustered enhancers (Chen et al. 2012), we next located the individual enhancers based on evidence of multiple TF binding in brain development using all publicly available transcription factor ChIP-seq data in cultured neural progenitors or embryonic cerebral cortex tissue (SRA and ENA dataset accession numbers provided in **Table S2**) (Estarás et al. 2012; Lodato et al. 2013; Webb et al. 2013; Lodato et al. 2014; Mateo et al. 2015; Sun et al. 2015; Sessa et al. 2017; Braccioli et al. 2018; Bertolini et al. 2019; Pagin et al. 2021). Aside from the *Sox2* CpG island promoter, regions that were approximately +3.7 kb (SRR2), +9.8 kb (chick N5 homologue; hereafter SRR10), +14 kb (adjacent to the chick N1 homologue; hereafter SRR14), and +17.1 kb (hereafter SRR17) downstream of the *Sox2* TSS showed binding of TFs associated with neuroepithelial/radial glia identity (ASCL1, BRN2, PAX6, SOX2), stem cell homeostasis (FOXO3, JUN, MAX, SOX9, TCF3), and neurogenesis (FEZF2, NFI, OLIG2, SMAD3, SOX4, SOX21, TBR1) (**Fig. 2A**; coordinates provided in **Table S3**). Notably, we observed only weak evidence of multiple TF binding at SRR1 (- 4.2 kb; chick N-2 homologue) in these datasets.

**Fig 2.**
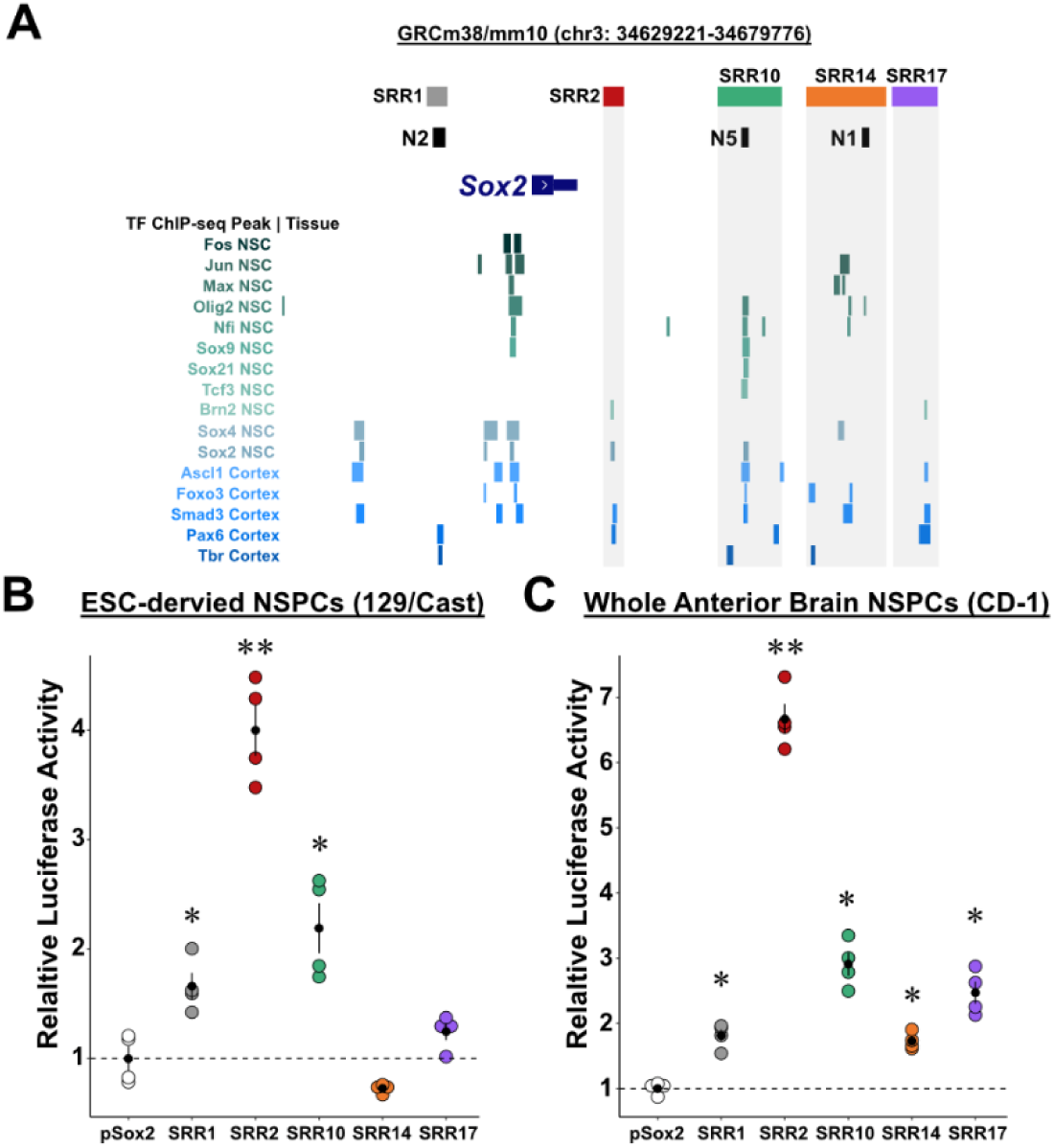
SRR2-18 contains multiple enhancers active in NSPCs from different sources. **(A)** The SRR2-18 genomic region is displayed on the UCSC Genome Browser (mm10). *Sox2* regulatory regions (SRRs, top) correspond to transcription factor-bound regions derived from NSPC and embryonic cerebral cortex ChIP-seq data sets compiled from public databases (bottom). **(B)** Clusters of transcription factor binding sites in neural cells identifying distinct elements capable of activating transcription in luciferase reporter assays using F1 embryonic stem cell (ESC) derived NSPCs. Dashed line indicates mean activity of the *Sox2* promoter (pSox2) alone. **(C)** The same luciferase reporter assays were performed in primary NSPCs generated from brain tissues dissected from mouse CD-1 embryos at E12.5. Error bars represent SEM. n ≥ 3. Significant differences from the empty pSox2 vector are indicated. (∗) P < 0.05, (∗∗) P < 0.01.

We next performed luciferase reporter assays to assess the sufficiency of each enhancer candidate to induce transcriptional activation. To account for regulatory bias potentially attributable to native enhancer-promoter compatibility (Martinez-Ara et al. 2022), we modified the reporter vector to replace the minimal promoter with the mouse *Sox2* promoter (-2209 to -347 bp of TSS) (**Supplemental Figure S2A**). Control experiments carried out in ESCs or NSPCs with a set of previously validated enhancers, namely SRR107 (a sub-module of the pluripotency-associated SCR) or the *Nes* intron 2 enhancer (Nes-I2) allowed us to verify that cell type-selective enhancer activity is reproduced with an ectopic reporter approach (Josephson et al. 1998; Zhou et al. 2014) (**Supplemental Fig. S2B**). F1 NSPCs transfected with reporter plasmids carrying SRR1, SRR2, or SRR10 had significantly increased reporter activity compared to cells with the *Sox2* promoter plasmid alone (*P* = 6.11 × 10^-3^, 2.13 × 10^-4^, or 7.3 × 10^-3^, Welch’s T Tests, **Fig. 2B**). To cross-validate the phenotype of F1 NSPCs in comparison to primary NSPCs established in the same EGF- and FGF2-containing culture medium, we generated NSPC lines from E12.5 whole brain tissue dissected rostral to the optic cup (including the telencephalon and the anterior diencephalon) from CD-1 mice. Despite the differences in genetic background, primary CD-1 NSPCs and F1 NSPCs show correlated (rho = 0.71, *P* = 3.04 × 10^-5^) activation of *Sox2* promoter-driven reporter gene expression from regulatory regions found upstream (SRR1) and downstream (SRR2-18) (**Fig. 2C**). These results suggest that SRR1 is an active enhancer in NSPCs derived from F1 ESCs and embryonic forebrain/midbrain tissue from a reference mouse strain, but that *Sox2* transcriptional regulation could be distributed across multiple TF-bound active downstream enhancers.

To test the role of this 16.7 kb downstream region independent of the upstream SRR1 region in neural lineage specification, we sought to decouple *Sox2* expression from SRR2-18 *cis*-regulation by CRISPR-Cas9 mediated deletion. We first generated an F1 reporter line with an mCherry insertion following the *Sox2* coding sequence and the cleavable P2A peptide on the *Mus musculus*^129^ allele to facilitate the isolation of clonogenic NSPCs for extensive deletion analyses. Homogeneity of mCherry reporter gene expression (>97%) in NSPCs differentiated from the *Sox2*-mCherry^129^ parent line was confirmed by flow cytometry (**Fig. 3A**). Accordingly, we used total RNA-sequencing (RNA-seq) to analyze the transcriptome of the bulk *Sox2*-tagged NSPCs. We found high similarity in the transcriptome of *Sox2*-mCherry^129^ NSPC replicates across different passage numbers in culture (rho > 0.8 on raw counts and rho > 0.98 on log-transformed counts, **Fig. 3B**). We compared the *Sox2*-mCherry^129^ NSPCs with other types of mouse neural progenitor cells using available datasets and hierarchical clustering analysis. These *Sox2*-tagged NSPCs grouped with Sox1^+^ neuroepithelial cells and Hes5^+^ telencephalic radial glia, suggesting that *Sox2*-mCherry^129^ NSPCs maintain consistent gene expression profiles that are similar to other well-characterized neural progenitor cell types (Bonev et al. 2017; Bertolini et al. 2019; Baumann et al. 2019) (**Fig. 3B**). Terminal differentiations of these NSPCs were carried out to verify astrocyte, oligodendrocyte, and neuronal lineage specialization competence. We followed a two-step protocol that involved cultivating *Sox2*-mCherry^129^ NSPCs and subclones in medium without exogenous epidermal growth factor (EGF) to facilitate exit from a self-renewing state with subsequent treatment with proprietary mixed lineage differentiation medium (NeuroCult™ Differentiation) for up to ten days (**Fig. 3C; Supplemental Fig. S3A**). As anticipated, the progenies differentiated from the Sox2-mCherry^129^ NSPCs showed decreased expression of cell cycle process genes (*Mki67*, *E2f2*), consistent with cell cycle exit, and induced expression genes associated with astrocyte (*Aldoc*, *Gfap*, *Sox9*), oligodendrocyte (*Cxcr4, Nkx2-2*), and neuronal (*Rbfox3/NeuN*, *Slc1a2*, *Sox21*) maturation (Sandberg et al. 2005; Hatada et al. 2008; Patel et al. 2010) (**Fig. 3D; Supplemental Fig. S3B**; **Table S4**). These findings indicate that the mCherry-tagged NSPCs are capable of gene expression changes associated with differentiation into neuronal and glial cell types.

**Fig 3.**
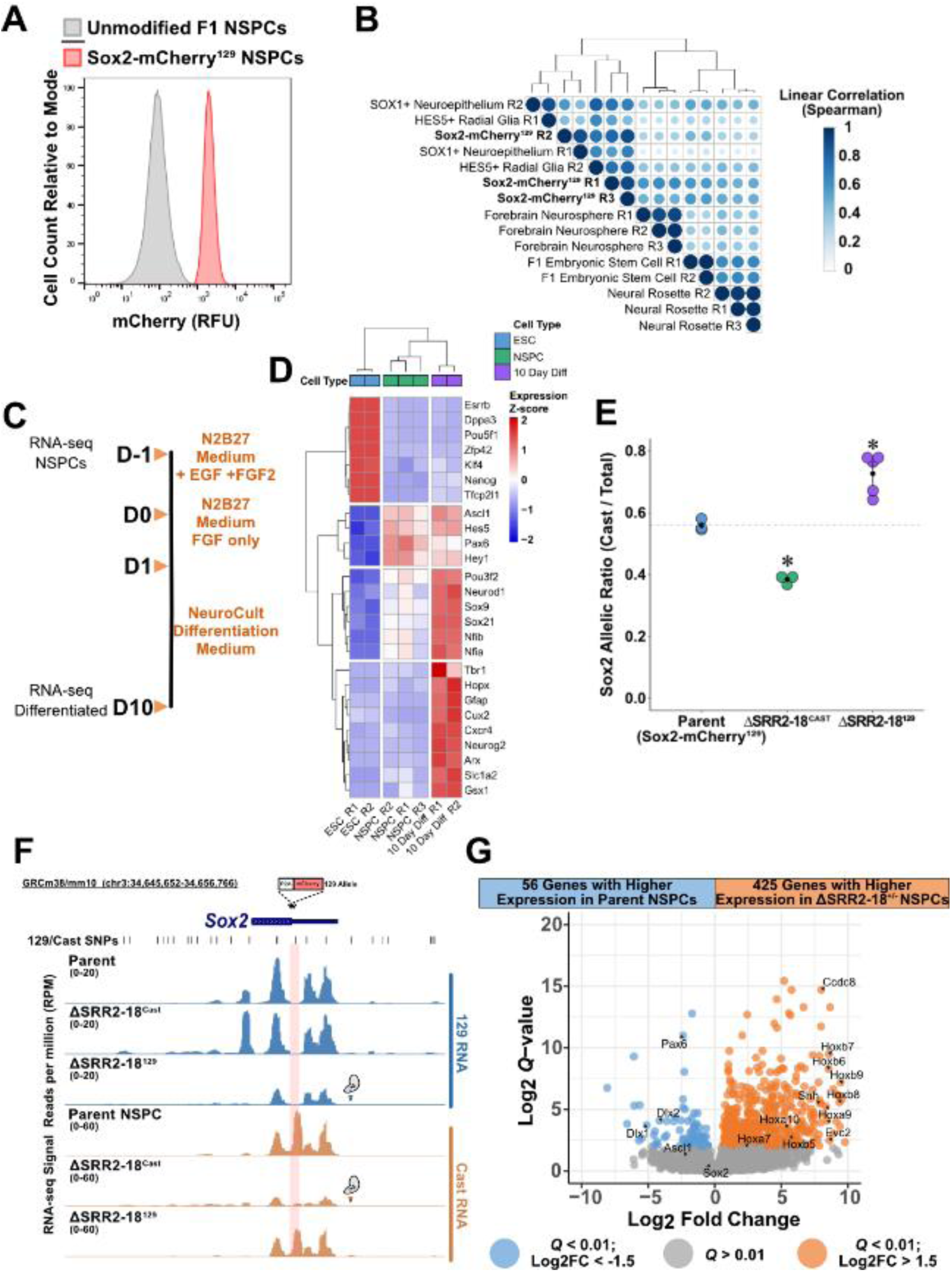
Monoallelic SRR2-18 deletion affects *Sox2* allelic dosage and positional identity of NSPCs. **(A)** Distribution of mCherry fluorescence in *Sox2*-mCherry^129^ NSPCs overlayed with genetically unmodified NSPCs. **(B)** Hierarchical clustering of sample correlations (Pearson rho represented by circle size and color) based on normalized transcript counts in bulk and transcription factor sorted subsets of neuroepithelial and neural progenitor cells from RNA-seq compiled from the GEO repository and this study. **(C)** Schematic of NSPC differentiation following sequential EGF and FGF2 withdrawal and successive undirected differentiation in a serum-containing supplement until differentiation day 10. **(D)** Hierarchical clustering of normalized counts across cell identity associated genes showing the patterns of gene expression changes between ESC-to-NSPC and NSPC-to-terminally differentiated cell transitions. **(E)** *Sox2* expression from the Cast allele in ESC-derived NSPCs by allele-specific RT-qPCR comparing the parent *Sox2*-mCherry^129^ lines to lines harboring monoallelic deletions of the SRR2-18 genomic region with the indicated genotypes. Expression levels for each allele are shown relative to the total transcript levels. Significant differences from the parent line values are indicated. (∗) P < 0.05. Error bars represent SEM. n ≥ 3. **(F)** Allele-sorted RNA-seq read pileup over the *Sox2* gene body, displayed on the UCSC Genome Browser (mm10). Red shaded bar indicates discriminatory 129/Cast single nucleotide polymorphism (SNP) on the 129 allele that is destroyed by the P2A-mCherry tag. Globular scissor protein icon denotes allele harboring a deletion of the SRR2-18 genomic region. **(G)** Volcano plot with the displayed number of differentially expressed genes between the parent and ΔSRR2-18^+/-^ NSPCs. Within table, number of genes that passed the cutoffs of absolute log2 fold-change > 1.5 and *Q* < 0.01. Q values represent the adjusted *Q* values computed with the Benjamini & Hochberg method for controlling false discovery rate (FDR). *P* values were extracted from a negative binomial regression model by Wald Chi-Squared tests.

From the *Sox2*-mCherry^129^ parent line, we generated numerous ΔSRR2-18 heterozygous monoclonal cell lines (three Cast and five 129 allelic deletions) to assess the *cis*-regulatory contribution of this enhancer cluster for *Sox2* expression and phenotype the deletion-harboring NSPCs. Allele-specific RT-qPCR analysis showed ΔSRR2-18^Cast^ NSPCs exhibited a significant decrease in the fraction of *Sox2* transcript from the Cast allele compared to NSPCs from the parent genetic background (*P* = 2.11 × 10^-4^, Welch’s T Test; **Fig. 3E**). Reciprocally, NSPCs with a ΔSRR2-18^129^ genotype displayed a significant increase in the Cast *Sox2* allelic ratio compared to the control representing the parent genetic background (*P* = 5.50 × 10^-3^, Welch’s T Test; **Fig. 3E**). These data indicate that the SRR2-18 region functions as a *cis*-regulator of *Sox2* in the neural lineage.

We next sought to investigate whether the monoallelic deletion of SRR2-18 specifically disturbs the allelic dosage of *Sox2* or if additional genes are also regulated in cis after this deletion. To account for mapping bias to reference mouse alleles, we assembled a SNP-substituted mouse genome using the 129S1/SvImJ sub-strain as a reference and N-masked discriminatory CAST/EiJ sub-strain SNPs (**Supplemental Fig. S3C**). We resolved the relative allelic expression at each locus to infer *cis* or *trans* modes of gene regulation. Profiles of allele-sorted RNA-seq reads showed that monoallelic deletion of SRR2-18 decreased steady state *Sox2* transcription from the targeted allele (**Fig. 3F**). Differential allele-resolved expression analysis in monoallelic deletants identified *Sox2* as the only chromosome 3 gene regulated by the SRR2-18 cluster (Cast/Total ratio log2FC > |0.5|; *Q* < 0.05; **Supplemental Fig. S3D**). These data show that SRR2-18 only regulates *Sox2* expression in a *cis*-acting mechanism. This analysis also highlighted that imbalances in allelic dosage between the 129 and Cast haplotypes are pervasive across a wide range of gene expression levels in F1 NSPCs, however these did not differ based on the targeted SRR2-18 allele.

Germline *Sox2* hypomorphism results in viable mice that present with posteromedial cerebral malformations and disrupted patterning of the anterior diencephalon (i.e. hypothalamus, pituitary) (Ferri et al. 2004; Kelberman et al. 2006; Langer et al. 2012). These models suggest that decreases in *Sox2* dosage below a certain threshold can negatively impact neural development. To test whether SRR2-18 mediated regulation of *Sox2* expression is required to establish the fundamental molecular properties of mouse NSPCs, we analyzed the total RNA-seq data in a manner agnostic to the hybrid cell haplotypes in ΔSRR2-18 heterozygous F1 NSPCs. Hereafter analyses of pooled ΔSRR2-18^129^ and ΔSRR2-18^Cast^ sample data is referred to as ΔSRR2-18^+/-^. Differential expression analysis revealed reproducible alterations to the ΔSRR2-18^+/-^ NSPC transcriptome, which included 425 upregulated and 56 downregulated RNAs (log2FC > |1.5|; *Q* < 0.01) (**Fig. 3G**). ΔSRR2-18^+/-^ NSPCs showed increased expression of the *Hoxb* locus genes that are associated with caudal patterning of the neural tube (*Hoxb6-9*), ventral patterning of the neural tube (*Shh*, *Sim2*, *Evc2*), and transmembrane proteins involved in cell migration and morphogenesis (*Icam1*, *Dnah5*, *Tmem179*) (**Fig. 3G; Table S5**). Parent Sox2-mCherry^129^ NSPCs had higher abundances of genes important for dorsal forebrain and midbrain development (*Pax6*, *Dlx2*, *Irx3*) (**Fig. 3G**) but also co-expressed genes associated with more caudal regions of the nervous system (*En2*, *Hoxa2*) (**Table S5**). Gene ontology (GO) analysis revealed terms with top enrichment scores (ES) among the upregulated gene set in ΔSRR2-18^+/-^ NSPCs, including, but not limited to, anterior/posterior pattern specification (*ES* = 0.69, *Q* = 3.1 × 10^-3^) and regionalization (*ES* = 0.61, *Q* = 7.5 × 10^-3^; **Supplemental Fig. S4A**; **Table S6**). Notably, ΔSRR2-18^+/-^ NSPCs show increased expression of *Wnt8a* which, along with *Wnt3a*, plays a niche-forming role in the development of caudal neural tissues (Ribes et al. 2009). In summary, NSPCs are sensitive to SRR2-18-driven *Sox2* transcriptional regulation, whereby ΔSRR2-18^+/-^ NSPCs activate caudal neural tube patterning pathways under steady-state culture conditions that normally maintain the self-renewal and identity NSPCs from more rostral brain regions.

### Biallelic deletion of SRR2-18 disrupts NSPC self-renewal and multipotency

To investigate the phenotype of NSPCs established without the SRR2-18 enhancer cluster, we derived monoclonal ESC lines with a biallelic SRR2-18 deletion (ΔSRR2-18^129/Cast^) and differentiated them into NSPCs. ΔSRR2-18^129/Cast^ NSPCs maintain Nestin and SOX2 immuno-reactivity while displaying reduced *Sox2* transcripts produced from both alleles across multiple clones (*P* = 2.3 × 10^-2^ for 129 allelic mutants and *P* = 6.7 × 10^-3^ for Cast allelic mutants, Dunn’s Test with B-H adjustment, **Fig. 4A, Supplemental Fig. S4B**). Since the reduction in *Sox2* transcript level in ΔSRR2-18^129/Cast^ NSPCs was only 42% compared to parent NSPC clones (**Fig. 4A**), we next sought to determine if the distribution SOX2 protein levels differed among parent, ΔSRR2-18^129^, and ΔSRR2-18^129/Cast^ NSPCs. Quantification of total intracellular SOX2 level by flow cytometry revealed that ΔSRR2-18^129/Cast^ NSPC cultures (40.0% decrease, *P* = 8.99 × 10^-7^, Welch’s T test), but not ΔSRR2-18^129^ heterozygous cells (10.2% decrease, *P* = 0.074, Welch’s T test), have fewer SOX2 expressing cells compared to the control line with the same genetic background (**Fig. 4B**). Similarly, ΔSRR2-18^129/Cast^ NSPC populations show a pronounced decrease in the mean SOX2 abundance compared to the parent line (33.7% decrease, P = 4.89 × 10^-5^, **Fig 4B**). Although SOX2 level tended to be lower in ΔSRR2-18^129^ NSPCs compared to the parent line, the effect was more subtle (15.0% decrease, P = 5.79 × 10^-2^). The observation that ΔSRR2-18^129^ heterozygous NSPCs have a comparable level of intracellular SOX2 to the parent line is surprising, but may be explained by compensatory post-transcriptional or post-translational mechanisms related to the caudal neural phenotype of these mutants at this developmental stage. To test whether the multi-modal SOX2 distribution observed in ΔSRR2-18^129/Cast^ NSPCs contributes to instability of the NSPC phenotype, we conducted growth curve analysis from multiple clones and different passage numbers. In maintenance culture conditions, ΔSRR2-18^129/Cast^ NSPCs initially show no difference in exponential-phase growth rate but adopt lower rates of cell division with prolonged time in culture (*P* < 7.8 × 10^-3^, Welch’s T tests, **Fig. 4C**), consistent with a self-renewal defect.

**Fig 4.**
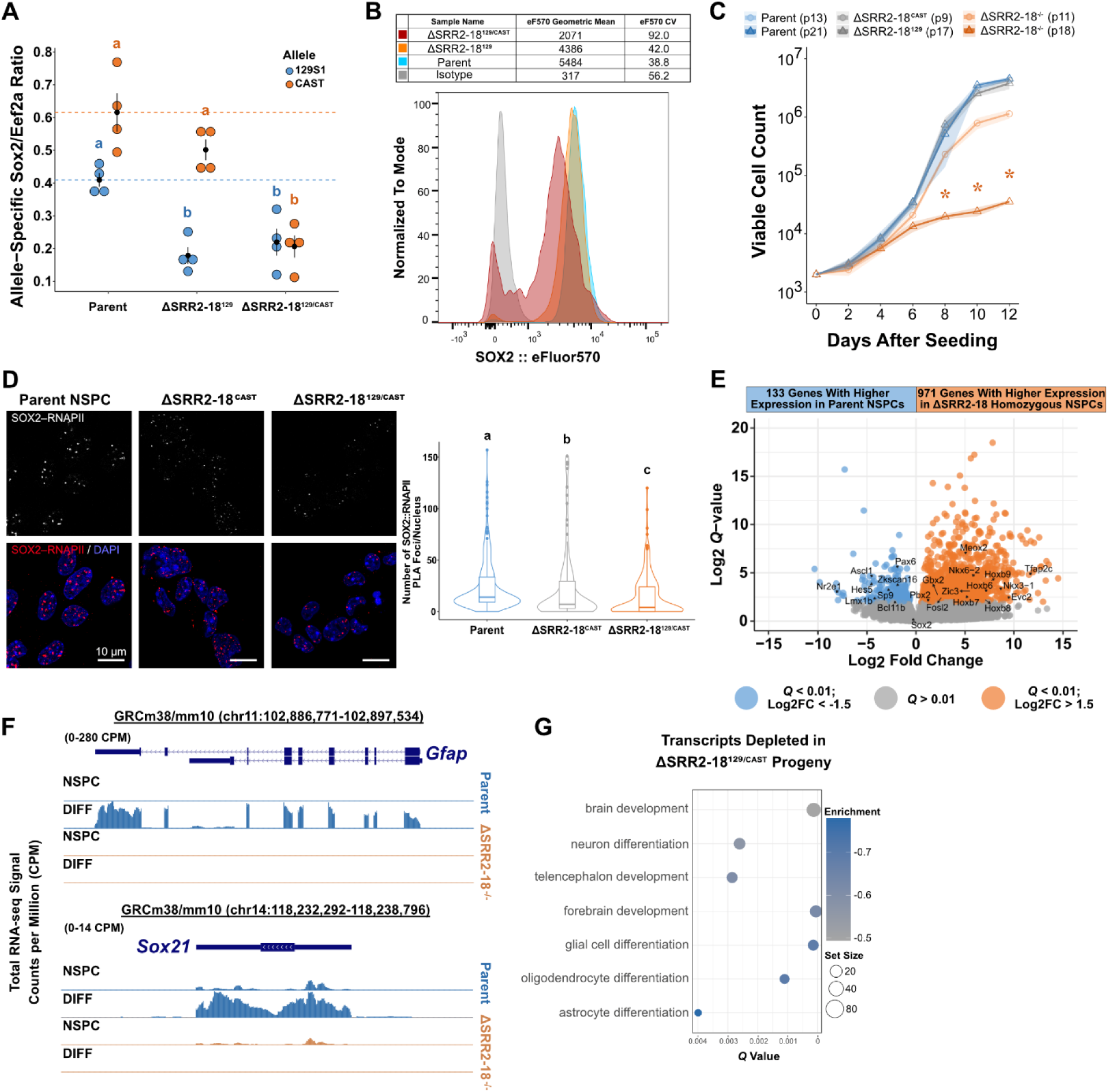
SRR2-18 is required for maintenance of the NSPC phenotype and differentiation *in vitro*. **(A)** *Sox2* expression in clonal NSPCs by allele-specific RT-qPCR comparing the indicated genotypes. Groups determined to be significantly different (P < 0.05) among multiple pairwise comparisons are labeled with different letters. To indicate P > 0.05, groups would be labeled with the same letter. Expression levels for each allele are shown relative to the reference transcript *Eef2a*. Error bars represent SEM. n ≥ 4. **(B)** Overlay histogram of flow cytometry data plotting the distribution of intracellular SOX2 level in parent, ΔSRR2-18^129^, and ΔSRR2-18^129/Cast^ NSPCs. The filled gray curve represents NSPCs stained with the isotype control conjugated to the same fluorophore which acts as a negative control. n=3. **(C)** Cell proliferation curves from different clonal NSPC lines and passage numbers showing differences in acute growth observed in ΔSRR2-18^129/Cast^ NSPCs. Groups determined to be significantly different (P < 0.05) among pairwise comparisons to the parent line are labeled with an asterisk (*). n ≥ 4 **(D)** Proximity ligation amplification (PLA) indicating the interaction between SOX2/RNAPII-S5P in parent, ΔSRR2-18^129^, and ΔSRR2-18^129/Cast^ NSPCs. Images show median-intensity Z stack projections. Scale bar distance is 50 µm. Box-and-whisker plot related to (D) enumerating PLA foci per nucleus. Boxes indicate interquartile range of intensity values and whiskers indicate the 10th and 90th percentiles. Outliers are represented by dots. Images were collected from at least three biological replicate samples and over 100 nuclei were quantified for each denoted genotype. Groups determined to be significantly different (P < 0.05) from one another are labeled with different letters. **(E)** Volcano plot with the displayed number of differentially expressed genes between the parent and ΔSRR2-18^129/Cast^ NSPCs that passed the cutoffs of absolute log2 fold-change > 1 and *Q* < 0.01. **(F)** Normalized RNA-seq tracks showing read pileup over the *Gfap* (astrocyte marker gene) and *Sox21* (neuronal maturation gene) loci, displayed on the UCSC Genome Browser (mm10). Signal tracks show greater transcript read coverage in libraries produced from the differentiated progeny (DIFF) of parent NSPCs to that of SRR2-18^129/Cast^ NSPC derivatives. **(G)** Bubble plot of the depleted gene sets of upregulated genes in differentiated progeny of ΔSRR2-18^129/Cast^ vs. control NSPCs generated by 10 days of *in vitro* differentiation. *Q* values represent the adjusted *P* values computed with the Benjamini & Hochberg method for controlling false discovery rate (FDR).

To obtain an accurate quantification of the functional relationship between SOX2 protein level and the frequency of transcriptional complex formation, we evaluated individual NSPCs by proximity ligation assays (PLA) with confocal microscopy (Dhaliwal et al. 2018; Dhaliwal et al. 2019). We quantified the protein interaction frequencies of SOX2 with RNA polymerase II (RNAPII) molecules engaged in the elongation of primary transcripts that are spatially resolved within single nuclei using an antibody targeting the phosphorylated serine 5 residue on the RNAPII carboxy-terminal domain (Komarnitsky et al. 2000). In the parent NSPCs, SOX2 interacts with RNAPII at a median of ∼ 21 visible nucleoplasmic foci (**Fig. 4D**). We observed that SOX2-RNAPII interactions in ΔSRR2-18^Cast^ NSPCs were significantly decreased compared to NSPCs from the parent genetic background (median = 7, *P* = 6.55×10^-6^, Wilcoxon Test; **Fig. 4D**). The frequency of SOX2-RNAPII association in NSPC nuclei were further decreased in the ΔSRR2-18^129/Cast^ line compared to that of monoallelic deletants (median = 4, *P* = 7.25×10^-3^, Wilcoxon Test; **Fig. 4D**). Together, these data suggest that deletion of SRR2-18 affects the participation of SOX2 protein in transcriptionally productive complexes through reduced *Sox2* transcription.

We also expanded our RNA-seq transcriptomic analysis to include these NSPCs established from homozygous ΔSRR2-18^129/Cast^ ESCs, which revealed ΔSRR2-18^129/Cast^ cells were even more dissimilar to the parent control line in a principal component analysis of RNA-seq data than heterozygous ΔSRR2-18^-/+^ NSPCs (**Supplemental Fig. S4C**). To assess if the gene expression profiles of NSPCs decoupled from SRR2-18 mediated regulation is distinct from complete Sox2 ablation, we compared our RNA-seq datasets to available transcriptome data from aggregates of self-renewing primary NSPCs (commonly referred to as neurospheres) established from wild-type and *Sox2*-ablated mice (Bertolini et al. 2019). Whereas *Sox2*^+/+^ neurospheres and our NSPCs from the *Sox2*-mCherry^129^ background grouped in a relatively small principal component space, the transcriptome of enhancer-deleted NSPCs markedly deviated from *Sox2*-knockout (KO) neurospheres (**Supplemental Fig. S4C**). Collectively, these findings suggest that regulatory perturbations of *Sox2* during neural differentiation leads to distinguishable transcriptomic signatures through the acquisition of alternative NSPC fates.

To identify transcript-level features that distinguish ΔSRR2-18^129/Cast^ NSPCs from the parent line, we performed differential gene expression analysis. A total of 133 transcripts showed a significant decrease in overall abundance and 791 transcripts showed a significant increase in biallelic SRR2-18 deletants (log2FC > |1.5|; *Q* < 0.01, Wald test under negative binomial model) (**Fig. 4E; Table S7**).

Annotation of the downregulated gene set in ΔSRR2-18^129/Cast^ NSPCs by gene ontology (GO) analysis revealed enriched terms, including, but not limited to, nucleobase biosynthesis (*ES* = -0.41, *Q* = 2.8 × 10^-2^) and CNS neuron differentiation (*ES* = -0.52, *Q* = 3.0 × 10^-2^, one-sided Fisher’s Tests) (**Supplemental Fig. S4D; Table S8**). Reciprocally, analysis of the induced gene set suggested that tube development (*ES* = 0.50, *Q* = 2.8 × 10^-3^) and positive regulation of MAPK cascade (*ES* = 0.57, *Q* = 1.3 × 10^-2^, one-sided Fisher’s Tests) were among the top enriched biological processes in ΔSRR2-18^129/Cast^ NSPCs (**Supplemental Fig. S4E**). These data indicated that biallelic SRR2-18-deleted cells showed dysregulated nucleobase metabolism and MAPK pathways as well as altered neurodevelopmental signatures.

To assess whether biallelic deletion of SRR2-18 perturbs further differentiation into post-mitotic CNS cell types, total RNA-sequencing was performed on bulk progenies differentiated from ΔSRR2-18^129/Cast^. Compared to differentiation of unmodified NSPCs, mixed lineage differentiation of ΔSRR2-18^129/Cast^ NSPCs displayed reduced expression of neuronal (*Sox21*, *Rbfox3*), astrocyte (*Gfap*, *Aldoc*) and early oligodendrocyte (*Cxcr4*) maturation associated genes (**Fig. 4F, Supplemental Fig. S5A**). The differentiated progenies of SRR2-18^129/Cast^ cells exhibited pronounced differences in the number of differentially expressed genes (1235 increased expression; 1133 decreased expression; log2FC > |1.5|; *Q* < 0.01, Wald test under negative binomial model) compared to differentiation progenies of parent NSPCs (**Supplemental Fig. S5B, Table S9**), indicating a profound defect in gene activation during differentiation. We observed a marked depletion of transcripts associated with biological processes involved in brain development, including, but not limited to, forebrain development (*ES* = -0.62, *Q* = 5.6 × 10^-5^, one-sided Fisher’s Test), telencephalon development (*ES* = -0.60, *Q* = 2.8 × 10^-3^, one-sided Fisher’s Test), and generation of neurons (*ES* = -0.31, *Q* = 7.4 × 10^-3^, one-sided Fisher’s Test) in differentiated cells from a ΔSRR2-18^129/Cast^ background compared to those in differentiated cells with intact SRR2-18 (**Fig. 4G; Table S10**). These results indicate that SRR2-18-mediated *Sox2* transcriptional regulation plays an important role in establishing a regional NSPC identity that supports differentiation into cell types found in the developing brain.

### Decoupling *Sox2* transcriptional regulation from SRR2-18 alters the accessible chromatin landscape of NSPCs

SOX2 has a well-recognized role in supporting a permissive chromatin environment for the proper induction of neuronal lineage genes during neurogenesis (Amador-Arjona et al. 2015; Bertolini et al. 2019). We set out to obtain a comprehensive understanding of the gene regulatory network elicited downstream from *Sox2* decoupled from SRR2-18 regulation. We performed genome-scale identification and analysis of protein-DNA interaction sites inferred from accessible chromatin regions using ATAC-seq. Reproducible ATAC-seq peaks were largely distributed at intergenic regions, annotated TSSs, and gene bodies (**Supplemental Fig. S6A-B**). Consistent with their transcriptional dysregulation of *Sox2*, ΔSRR2-18^129/Cast^ NSPCs showed a decrease in chromatin accessibility at the *Sox2* promoter compared to that in parental cells (**Fig. 5A**). Analysis of ATAC-seq reads within the consensus peak set identified 88,910 regions with increased and 65,152 regions with decreased chromatin accessibility (log2FC > |1.5|; *Q* < 0.00001, Wald test under negative binomial model) (**Fig. 5B**; **Table S11**). These data suggest that *Sox2* enhancer-perturbed NSPCs adopt a distinct regulome that reflects the unique spatial and functional identity of ΔSRR2-18^129/Cast^ cells.

**Fig 5.**
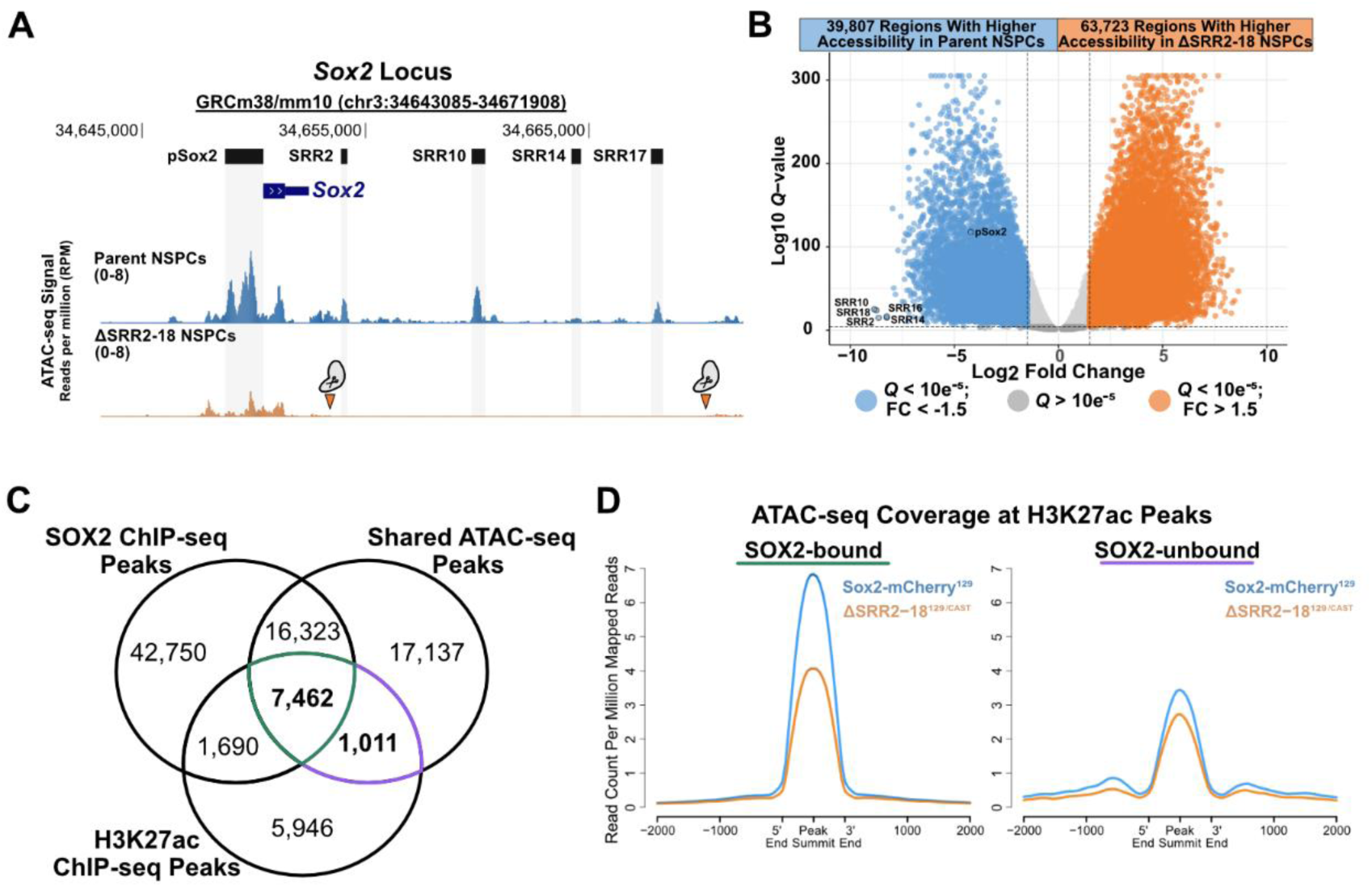
Deletion of SRR2-18 alters the accessible chromatin landscape of NSPCs. **(A)** The *Sox2* gene and SRR2-18 genomic regions displayed on the UCSC Genome Browser (mm10). *Sox2* promoter and regulatory regions (SRRs, top) in NPSCs and ATAC-seq tracks from parent and SRR2-18^129/Cast^ NSPCs (bottom). Globular scissor protein icon denotes the margins of a biallelic SRR2-18 deletion. **(B)** Volcano plot, with number of differentially accessible genomic regions between the parent and ΔSRR2-18^129/Cast^ NSPCs. Within table, number of regions that passed the cutoffs of absolute log2 fold-change > 1.5 and *Q* < 0.00001. *Q* values represent the adjusted *P* values computed with the Benjamini & Hochberg method for controlling false discovery rate (FDR). *P* values extracted from a negative binomial regression model by Wald Chi-Squared tests in DiffBind. **(C)** Shared accessible genomic regions between parent and ΔSRR2-18^-/-^ NSPCs divided into SOX2-bound, SOX2-unbound, H3K27ac-modified and non-H3K27ac modified subsets based on NSPC ChIP-seq data available in the GEO data depository. **(D)** Normalized ATAC-seq read coverage at SOX2-bound/H3K27ac and SOX2-unbound/H3K27ac accessible genomic regions between parent and ΔSRR2-18^129/Cast^ NSPCs.

We considered that those candidate regulatory elements represented in the set of decreased chromatin accessibility regions in ΔSRR2-18^129/Cast^ NSPCs were potential SOX2 targets in multipotent neural progenitors. To this end, we intersected our consensus ATAC-seq peak set shared between parent and deletant NSPCs with publicly available H3K27ac ChIP-seq peaks from Sox1-GFP^+^ *in vitro* differentiated NSPCs (Bonev et al. 2017). We reanalyzed ChIP-seq data obtained from GEO to establish a non-redundant set of SOX2-occupied regions in NSPCs (74,233 unique regions; **Table S12**). This allowed us to distinguish between regions of accessible chromatin associated with SOX2 regulation and loci with a similar chromatin profile but that are not direct SOX2 targets (**Fig. 5C**). Average profiles of ATAC-seq signal in the Sox2-mCherry^129^ NSPCs across qualifying genomic intervals showed enrichment of ATAC-seq signal at SOX2-bound regions (mean summit coverage = 6.7 RPM at 7,462 retained peaks) compared to signal at non-SOX2 target H3K27ac-modified regions (mean summit coverage = 3.2 RPM at 1,011 retained peaks) (**Fig. 5D**). As expected for an accessible and acetylated chromatin state, motif enrichment analysis of SOX2-bound regions predominantly recovered SOX, E26 transformation specific (ETS), and activator protein-1 (AP-1) motifs corresponding to the transcription factor families previously recognized to contribute to NSPC self-renewal (**Table S13**) (Pagin et al. 2021). Furthermore, SOX2-bound and H3K27ac-modified regions display a lower degree of ATAC-seq signal enrichment in ΔSRR2-18^129/Cast^ cells compared to that in parent NSPCs (**Fig. 5D**), which could reflect the reduction in SOX2 protein levels in these cells.

To better understand the biological processes likely to be affected by altered protein-chromatin interactions genome-wide, GO analysis was performed on the differentially accessible regions annotated to the nearest TSS. Terms overrepresented in the set of ATAC-seq peaks showing decreased accessibility in ΔSRR2-18^129/Cast^ NSPCs were related, but not limited to, forebrain development (*ES* = -0.34, *Q* = 2.9×10^-10^, one-sided Fisher’s Test) and neurogenesis (*ES* = -0.33, *Q* = 1.7×10^-35^, one-sided Fisher’s Test) (**Fig. 6A; Table S14**). Integrative analysis of the regulome and transcriptome in ΔSRR2-18^129/Cast^ NSPCs revealed that multiple proneural factors known to prime and regulate neuronal specification show a loss of transcriptional activation and chromatin accessibility, including Achaete-scute family TF 1 (*Ascl1*; also known as *Mash1*) (**Fig. 6B**). Reciprocally, genes near ATAC-seq peak regions with increased chromatin accessibility in ΔSRR2-18^129/Cast^ NSPCs included, but were not limited to, terms associated with regulation of anatomical structure morphogenesis (*ES* = 0.37, *Q* = 8.4×10^-38^, one-sided Fisher’s Test), anterior/posterior pattern specification (*ES* = 0.31, *Q* = 3.0×10^-8^, one-sided Fisher’s Test), and neural tube development (*ES* = 0.25, *Q* = 4.2×10^-4^, one-sided Fisher’s Test) (**Fig. 6C; Table S15**). We observed a specific increase in chromatin accessibility and transcription in *Hoxb* locus genes (*Hoxb5-9*) related to caudal neural tube patterning (**Supplemental Fig. S6C**), consistent with the regional identity bias of SRR2-18-deleted NSPCs observed by RNA-seq. In the developing vertebrate trunk, caudal type homeobox (CDX) factors function upstream of the *Hox* loci and are required for spinal cord specialization and patterning (Chawengsaksophak et al. 2004; Mazzoni et al. 2013). ΔSRR2-18^129/Cast^ NSPCs exhibit a switch-like gain of active transcriptional features at the *Cdx2* locus compared to the parent Sox2-mCherry^129^ line (**Fig. 6D**). These findings indicate that the induced expression of caudal positional factors and downregulation of factors involved in neuronal cell commitment observed in ΔSRR2-18^129/Cast^ NSPCs are supported by genome-wide alterations in the accessible chromatin landscape.

**Fig 6.**
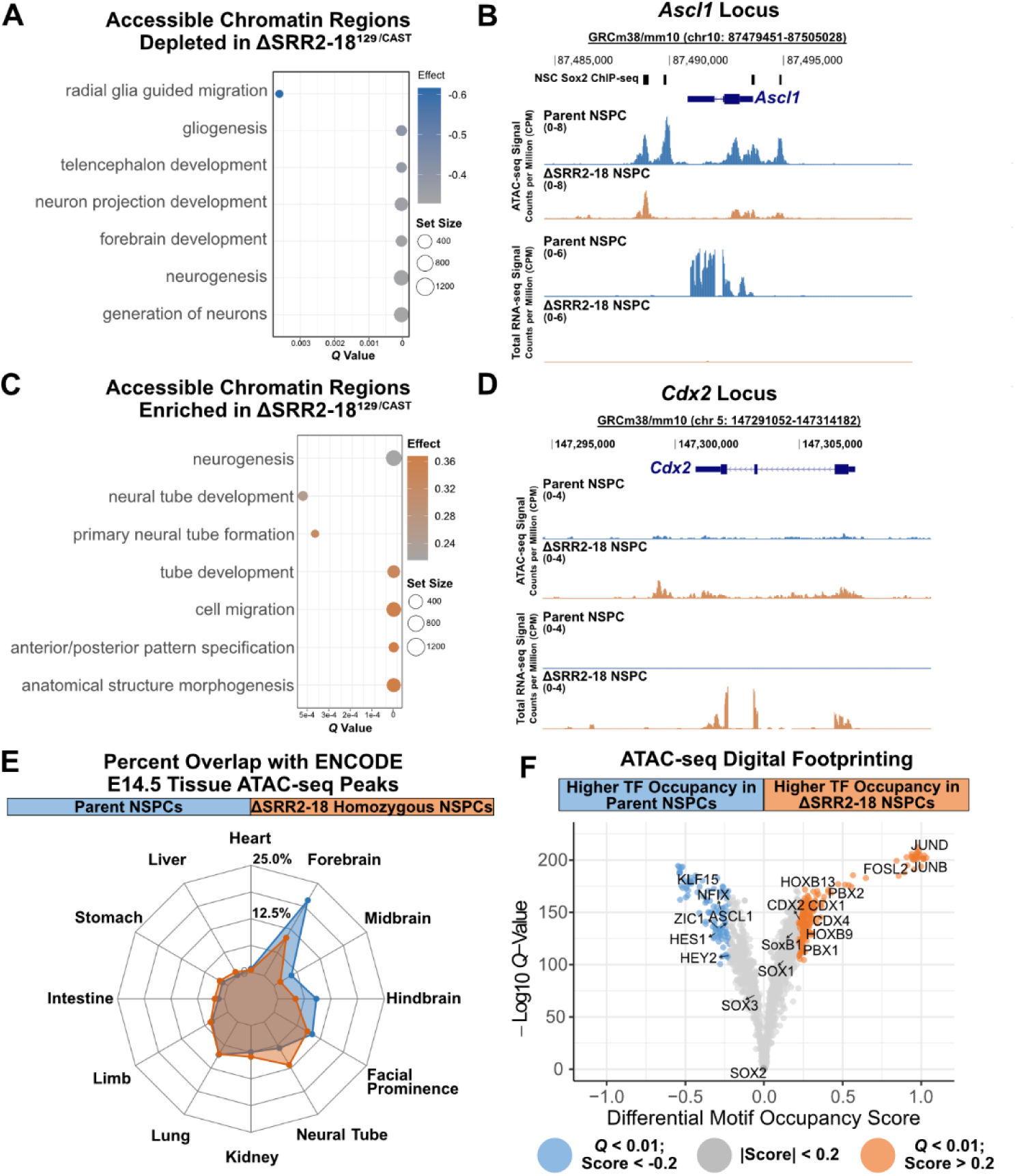
Induction of caudal patterning factors and chromatin remodeling in NSPCs decoupled from SRR2-18 mediated regulation. (A) Bubble plot of the underrepresented genomic regions in the set of decreased accessibility regions in ΔSRR2-18^129/Cast^ vs. control NSPCs. (B) Normalized ATAC-seq and RNA-seq tracks over the *Ascl1/Mash1* gene and flanking genomic regions displayed on the UCSC Genome Browser (mm10). Signal tracks show reduced accessibility at the *Ascl1* promoter and decreased total *Ascl1* RNA in ΔSRR2-18^129/Cast^ NSPCs. (C) Bubble plot displaying enriched genomic regions in the set of increased accessibility regions in ΔSRR2-18^129/Cast^ vs. control NSPCs. (D) Normalized ATAC-seq and RNA-seq tracks over the *Cdx2* gene and flanking genomic regions displayed on the UCSC Genome Browser (mm10). Signal tracks show increased accessibility at the *Cdx2* locus and increased total *Cdx2* RNA in ΔSRR2-18^129/Cast^ NSPCs. (E) Radar diagram of accessible genomic regions called from ATAC-seq data in parent NSPCs (blue border) and SRR2-18^129/Cast^ NSPCs (orange border) overlapping ENCODE consensus ATAC-seq peaks from E14.5 tissue samples. Height of the borders at each tissue intersection corresponds to the percentage of non-promoter accessible genomic regions that overlap. (F) Volcano plot, with the differential motif occupancy scores reflecting global changes in transcription factor binding in the consensus ATAC-seq peakset between the parent and ΔSRR2-18^129/Cast^ NSPCs from TOBIAS footprinting analysis. *Q* values represent the adjusted *P* values computed with the Benjamini & Hochberg method for controlling false discovery rate (FDR).

We explored the possibility that biallelic SRR2-18 deletion activates gene regulatory programs associated with posterior embryonic cell types. To assess the similarity of ΔSRR2-18^129/Cast^ NSPCs with tissue-specific chromatin accessibility signatures, we intersected our NSPC ATAC-seq peaks with a collection of embryonic day 14.5 (E14.5) excised mouse tissue ATAC-seq peaks from the ENCODE Consortium (**Table S1**). *Sox2*-mCherry^129^ ESC-derived NSPCs show the greatest overlap in accessible chromatin intervals with forebrain ATAC-seq peaks (non-promoter overlaps = 41,737, odds ratio = 114.8; **Fig. 6E**), consistent with extensive neurogenesis in the forebrain at this developmental stage.

Overlap with other anterior neural plate-derived tissues – the midbrain and hindbrain – is consistent with control NSPCs exhibiting a neural chromatin landscape configuration without a well-defined ventral-dorsal identity (non-promoter overlaps = 16,630 & 22,093, odds ratio = 98.5 & 127.1; **Fig. 6E**). Parent NSPCs exhibited fewer accessible chromatin interval intersections with the caudal portion of the neural tube (non-promoter overlaps = 21,118, odds ratio = 28.4; **Fig. 6E**). Indeed, reproducible ATAC-seq peaks from parent NSPCs were significantly enriched in the forebrain peak set (odds ratio = 15.0, *P* = 1.28 × 10^-35^, hypergeometric test) but not in the neural tube (odds ratio = 0.3, *P* = 1, hypergeometric test). The ΔSRR2-18^129/Cast^ NSPC were observed to have fewer shared accessible chromatin regions with forebrain tissue (non-promoter overlaps = 26,672, odds ratio = 18.1) and a relatively greater degree of overlap with neural tube tissue (non-promoter overlaps = 27,146, odds ratio = 92.9; **Fig. 6E**). Accessible chromatin regions from ΔSRR2-18^129/Cast^ NSPC were not significantly enriched in forebrain peaks (odds ratio = 1.7, *P* = 1, hypergeometric test) but did significantly overlap with the neural tube peak set (odds ratio = 2.9, *P* = 7.21 × 10^-11^, hypergeometric test).

We performed hierarchical clustering on normalized ATAC-seq reads we reanalyzed from open access data on samples isolated from brain and spinal cord neural tissues (**Table 2**) (Xu et al. 2018; Lattke et al. 2021; Delás et al. 2023). Cell type-specific patterns in chromatin accessibility with appropriate biological process enrichment could be identified (**Supplemental Fig. S7A-B**); however, the majority of ΔSRR2-18^129/Cast^ NSPC open chromatin regions form a partially overlapping cluster with moderately accessible genomic regions in ventral neural tube progenitors (Cluster 7). GO terms from the nearest genes to cluster 7 regions are associated with chordate development and cell differentiation in the spinal cord (**Supplemental Fig. S7A-B**). Predictably, these bulk cell and tissue analyses fell short of pinpointing a specific developmental cell state associated with the accessible chromatin profile of ΔSRR2-18^129/Cast^ NSPCs (**Supplemental Fig. S7A-C**).

To assess how genome-wide protein-DNA interactions are altered in ΔSRR2-18^129/Cast^ NSPCs, we next identified the repertoire of TF motifs found at accessible chromatin regions. For this, we carried out digital footprinting, an approach that infers TF occupancy from ATAC-seq data based on the transposition frequency at base-pair resolution, after estimation and correction of positional bias (Bentsen et al. 2020). We found several dozen sequence-specific TFs with a differential occupancy score outside of statistical thresholds in NSPCs between the two genotypes (score > |0.2|, *Q* < 0.01; **Fig. 6F**, Two-sided one-sample T test under background distribution). Notably, SOX2 footprints were not differentially enriched between parental and ΔSRR2-18^129/Cast^ NSPCs which may be due to remaining SOX2 protein in these cells; however, a relative lack of motif content differences across SoxB1 family members could not be ruled out. Moreover, functional compensation between co-expressed SoxB1 family transcription factors in neural tissues is well documented (Graham et al. 2003; Miyagi et al. 2008). Several transcription factors that regulate NSPC proliferation and neuronal differentiation in various parts of the brain were markedly depleted in ΔSRR2-18^129/Cast^ NSPCs, including KLF15, NFIX, ASCL1, and the Notch downstream effectors HES1 and HEY2 (**Fig. 6F**).

Footprinting analyses highlighted several TF motifs that participate in the AP-1 complex and are enriched within NSPCs derived without SRR2-18 mediated *Sox2* regulation (**Fig. 6F**). Additionally, spinal cord specification and patterning factors such as CDX2/4, HOXB, and PBX1/2 also contribute to the altered chromatin accessibility profile in ΔSRR2-18^129/Cast^ NSPCs (**Fig. 6F**). These results indicate that decoupling Sox2 expression from the SRR2-18 enhancer cluster induces a caudal region-specific identity. This loss is characteristic of cultured NSPCs driven by posterior regionalization factors in a regulatory network that is sensitive to *Sox2* transcript abundance and protein function.

## Discussion

The mechanisms of enhancer-mediated lineage- and cell type-specific gene activation at different developmental stages or spatial contexts are unclear. Previous studies have suggested that enhancer switching is common during cell differentiation by comparing lineage-specific TF binding sites in stem/progenitor and mature cells (Soleimani et al. 2012; Alder et al. 2014; A.F. Chen et al. 2018). We found that *Sox2*, a key factor in pluripotency and NSPC fate (Masui et al. 2007; Foshay and Gallicano 2008), exhibited a dynamic temporal pattern of *Sox2* RNA levels during mouse ESC to neural differentiation. Leveraging the genetic background of *M. musculus*/129 × *M. castaneus* ESCs and their derivatives, we demonstrated that these dynamics were coincident with a switch in regulatory control of *Sox2* from the pluripotency associated SCR to a proximal downstream region that determines *Sox2* transcription during the ESC-to-NSPC transition. Additionally, we found that the SRR2-18 region is necessary to maintain the level of *Sox2* transcriptional output in ESC-derived NSPCs under self-renewing conditions. Using an allelic deletion series, we showed that SRR2-18-mediated *Sox2* regulation is critical for sustaining the proliferative rate of NSPCs, establishing a region-specific neural progenitor identity, and activating neuronal differentiation genes. We identified patterns in differential chromatin accessibility and TF motif representation between parental and ΔSRR2-18^129/Cast^ NSPCs that propose how neural *Sox2* regulation safeguards a neurogenic program and a regionally biased neural progenitor identity through gene regulatory network activation.

Other investigations have described the regional selectivity of enhancer-mediated gene expression for candidate *Sox2* proximal enhancers in neural tissues using transgenic chick or mouse reporter assays (Uchikawa and Kondoh 2016). The proximal enhancers SRR1 and SRR2 show reporter gene activity in neurospheres and spatially restrict expression to the telencephalon in murine neural development (Zappone et al. 2000; Miyagi et al. 2004; Miyagi et al. 2006), however, neither SRR1 nor candidate distal enhancers (Beagan et al. 2017; Chakraborty et al. 2023) are sufficient to functionally compensate for the loss of SRR2-18-mediated regulation of *Sox2* in F1 NSPCs. To this point, we observed no difference in the level of *Sox2* knockdown in compound monoallelic ΔSRR1;ΔSRR2-18 (31-42%) and monoallelic ΔSRR2-18 NSPCs (32-38%). A similar level of *Sox2* transcriptional deficiency in NSPCs of the developing forebrain has been described using a *Sox2* haplodeficient mouse model harboring a heterozygous deletion of SRR1 on the intact copy of *Sox2* (Ferri et al. 2004). However, in this case, reduced *Sox2* level in NSPCs *in vivo* was dependent on engineering a null *Sox2* allele. These findings indicate that, despite its functional sufficiency to activate gene transcription in primitive neural cell populations, SRR1 is dispensable for *Sox2* expression in a mouse NSPC model expressing *Sox2* biallelically.

Conditional biallelic deletion of the *Sox2* coding sequence resulted in impaired hippocampal NSPC survival and differentiation competence, as well as self-renewal defects in primary NSPCs upon extended maintenance in culture (Favaro et al. 2009; Bertolini et al. 2019). These studies highlighted the roles of immediate early genes and cytokine signaling in the transcriptional amplification cascades linked to NSPC self-renewal. Because clustered enhancers may function in a partially redundant manner within the genomic context (Moorthy et al. 2017), our approach was to delete the 16.7 kb region constituting the clustered *Sox2* proximal elements. Using allele-specific transcriptomics, we confirmed that, aside from *Sox2*, no other coding or non-coding RNAs were controlled in a direct *cis*-acting mechanism attributable to SRR2-18. Decoupling *Sox2* regulation from the SRR2-18 enhancer cluster was permissive to neural induction; however, NSPCs harboring a SRR2-18 deletion on even one allele showed increased expression at multiple positional factor genes that specify the caudal identity of neural cells. The organizing stimulus of gene expression programs characteristic of caudal neural cell fates in adherent NSPCs is unclear; however, the increased expression of *Wnt8a* observed in mutant NSPCs could, if secreted into the culture microenvironment in a paracrine manner, promote the maintenance of caudally fated NSPCs. A reduction in *Sox2* dosage during axial regionalization *in vivo* has been reported to preconfigure the regulatory landscape of the caudal epiblast for WNT signaling input to specify posterior neural fates (Metzis et al. 2018; Blassberg et al. 2022). Additionally, fibroblast growth factor (FGF) signaling is involved in multiple stages of neural tube development, including specification of spinal cord identity, caudal extension and patterning, and directing of the onset of ventral neural tube identity and patterning (Reviewed in (Diez Del Corral and Morales 2017)). However, given that all NSPC culturing media (containing exogenous FGF2 and EGF) are chemically defined and devoid of exogenous retinoids and Smoothened receptor ligands, the patterning responses observed in ΔSRR2-18^129/Cast^ NSPCs are consistent with cell-derived instructive factors. It will be interesting to examine if SRR2-18-mediated SOX2 regulation can acutely modify embryonic cell responses to FGF or WNT signaling during neural induction and axial regionalization.

Regulating chromatin accessibility by the cumulative action of protein binding and chromatin modification is an important mechanism for controlling gene expression patterns (Thurman et al. 2012). Changes in chromatin accessibility or local genome topology can alter gene regulation programs during cell differentiation, leading to the establishment of unique spatial and functional cell identities (Iwafuchi-Doi et al. 2016). SOX2 regulates the transcriptional permissiveness of its target genomic sites in multiple cellular contexts and facilitates regulatory chromatin-chromatin interactions in NSPCs that safeguard against depletion of the progenitor cell pool during brain development (Bertolini et al. 2019; Dodonova et al. 2020). In brain-derived neural progenitors, SOX2 transcriptionally primes a set of conserved basic helix-loop-helix TFs known as proneural genes, which induce neuron differentiation during brain development (Amador-Arjona et al. 2015). In this study, ESC-derived NSPCs exhibited transcriptome and accessible genome features similar to forebrain progenitors and showed primarily anterior brain tissue chromatin accessibility profiles. We observed specific patterns in genome-wide chromatin accessibility in ΔSRR2-18^129/Cast^ cells, which are associated with footprints for TFs that have previously established roles in rostral-caudal regionalization of the nervous system. To our surprise, the highest scoring differential motif occupancy scores were classical immediate-early genes of the JUN, FOS, and ATF protein families. Components of the AP-1 complex, these multifunctional TFs play roles in cell proliferation, cell migration, neuronal maturation and activity, and the cell stress response (Reviewed in (Bejjani et al. 2019)). Different genetically engineered mouse models deficient for stress- or mitogen-activated phosphorylases upstream of the AP-1 complex show defects in neural tube closure (Sabapathy et al. 1999; Chi et al. 2005); however, there are currently no high-confidence AP-1 TF family targets implicated in neural tube morphogenesis. SOX2 cooperates with distinct TF repertoires to target diverse genomic sites across different *Sox2*-expressing developmental fates, including in the spinal cord (Hagey et al. 2018). Given that ESC-derived NSPCs were propagated in adherent culture and FGF2-containing medium prior to transcriptome and regulome profiling, the temporal progression in regulatory mechanisms that direct this shift in target DNA recognition remains to be determined.

Elucidating the regulatory mechanisms of *Sox2* is essential for understanding the transcriptional programs controlling neurodevelopment and the spatial diversity of NSPC fates. In this study, we examined the *Sox2* locus and identified a proximal enhancer cluster that regulates *Sox2* transcription NSPCs. Using ectopic reporter assays and allele-specific deletion analyses, we determined that the SRR2-18 enhancer cluster functions as a *cis*-regulator of *Sox2* in NSPCs. We show that SRR2-18-driven *Sox2* allelic dosage is necessary for establishing the rostro-caudal identity of ESC-generated NSPCs.

## Materials and Methods

### Embryonic Stem Cell Culture

F1 (*M. musculus*/129 × *M. castaneus*) ESCs were obtained from Barbara Panning, University of California San Francisco (Mlynarczyk-Evans et al. 2006). Cells were maintained on 0.1% (mass/vol) porcine gelatin coated plates and Dulbecco’s Modified Eagle Medium (Thermo Fisher 11960044) supplemented with 15% (vol/vol) ESC-qualified fetal bovine serum (Wisent Bioproducts 920040), 0.1 mM nonessential amino acids (Thermo Fisher 11140050), 1 mM sodium pyruvate (Thermo Fisher 11360070), 2 mM GlutaMAX (Thermo Fisher 35050061), 0.1 mM 2-mercaptoethanol (Thermo Fisher 21985023), 1X Penicillin-Streptomycin (Thermo Fisher 15140122), 1000 U/mL recombinant leukemia inhibitory factor (LIF), 3 μM GSK3β inhibitor CHIR99021 (Millipore Sigma 361559-5MG), and 1 μM MEK inhibitor PD0325901 (Millipore Sigma 444966-5MG), referred to here as “LIF/2i medium”. Incubators were humidified and maintained at 37°C, 5% CO_2_ under normoxic conditions. All cultures were confirmed to be free of mycoplasma contamination during routine testing with the Mycoplasma PCR Detection Kit (BioVision Inc. K1476-100).

### *In Vitro* Neural Differentiation and Neural Stem/Progenitor Cell Culture

F1 mouse ESCs were differentiated toward NSPC *in vitro* according to established protocols (Ying et al. 2003). Briefly, ESCs were seeded into 10 cm gelatinized culture plates in LIF/2i medium (day 0) at a density of 1 × 10^4^ cells per cm^2^. On day 1, culture medium was changed to a 1:1 mixture of Neurobasal medium (Thermo Fisher 21103049) and DMEM/F12 medium (Thermo Fisher 10565018) supplemented with 1X N2 (Thermo Fisher 17502048), 1X B27 without Vitamin A (Thermo Fisher 12587010), 2 mM GlutaMAX, 10 mM HEPES (Thermo Fisher 15630080), 0.015% Fraction V BSA (mass/vol), 0.1 mM 2-mercaptoethanol and 1X Penicillin-Streptomycin, referred to here as “N2B27 medium”. On day 5, adherent differentiating cells were detached with Accutase (Millipore Sigma A6964-500ML) and transferred to low adherence 10 cm plates in N2B27 medium supplemented with 10 ng/mL murine epidermal growth factor (Peprotech Inc. 315-09-100UG) and 10 ng/mL fibroblast growth factor 2 (Cedarlane Labs CLCYT386), referred to here as “NSE medium”. On days 10-12, the cell aggregates formed in suspension culture were dissociated with Accutase and seeded onto T25 flasks coated with 100 µg/mL of poly-D-Lysine (PDL) (Millipore Sigma 27964-99-4) and 10 µg/mL laminin (Millipore Sigma L2020) in NSE medium at a density of 2 × 10^4^ cells per cm^2^. This is considered passage 0 (p0) of neural stem/progenitor cell cultures. NSPCs were seeded into poly-D-lysine- and laminin-coated T25 flasks in NSE medium at a density of 2 × 10^4^ cells per cm^2^ and passaged every 4-5 days for a maximum of 25 passages. NSPC multipotency was assessed by differentiation into neuronal, astrocyte, and oligodendrocyte lineages following removal of EGF from NSE medium for one day and later by culture in NeuroCult™ Differentiation Medium (Stem Cell Technologies 05704) for an additional 9 days.

### CRISPR-Cas9 Mediated Deletion and Fluorescent Reporter Line Generation

Cas9-mediated deletions were carried out as previously described (Moorthy and Mitchell 2016). Briefly, Cas9 targeting guide RNAs (gRNAs) flanking predicted enhancer regions are provided in **Table 16**. gRNA plasmids were assembled in the gRNA Cloning Vector (a gift from George Church; Addgene plasmid # 41824), using the protocol described by Mali *et al*. (Mali et al. 2013). The sequence of the resulting guide plasmid was confirmed by Sanger sequencing. For deletions, F1 ESC were transfected with 5 μg each of 5′ gRNA, 3′ gRNA, and pCas9_GFP plasmid (a gift from Kiran Musunuru; Addgene plasmid # 44719) (Ding et al. 2013). To tag *Sox2* at the C-terminal, we constructed a homology directed repair donor template using Gibson assembly. The following sequences were assembled using the Gibson Assembly cloning kit (New England Biolabs E5510S): mCherry2-N1 amplicon (a gift from Michael Davidson; Addgene plasmid # 54517), P2A sequence through oligos, and homology arms amplified from F1 genomic DNA excluding the gRNA target site in the 3’ arm (**Table 17**). For homology-directed repair knock-in, F1 ESC were transfected with 5 μg each of *Sox2* 3’ untranslated region (UTR) gRNA and pCas9_GFP plasmid, as well as 10 μg donor fragment purified from EcoRV restriction digest. Transfections were performed using the Neon™ Transfection System (Thermo Fisher Scientific). Forty-eight hours after transfection, cells were sorted on a BD FACSAria for GFP-positive cells (for deletion experiments) or mCherry-positive cells (for tag knock-in). The collected cells were seeded at 1 × 10^4^ per cm^2^ on 10 cm gelatinized culture dishes and grown for 5–6 days until large well-defined colonies formed. Isolated colonies were picked and propagated for genotyping and gene expression analysis. For DNA extraction from 96-well plates, cells were washed once with PBS and subsequently lysed in a humidified chamber with 50 µL 1X cell lysis buffer using the prepGEM Universal Kit (PUN0100, MicroGEM), according to manufacturer’s recommendations. The 96-well plates were screened for deletions using allelic-specific qPCR-based assays. All deletions were confirmed by Sanger sequencing and analysis using primers 5′ and 3′ from the gRNA target sites. SNPs within the amplified product confirmed the genotype of the deleted allele.

### Derivation of Primary NSPCs from Mouse Embryonic Brain Tissue

All animal experiments were approved under animal use permit #20011749 by the University Animal Care Committee (UACC) of the University of Toronto. Embryo collections from wildtype CD-1 (CD1(ICR) from Charles Rivers) were made using timed mating. On E12.5, pregnant CD-1 females were euthanized by cervical dislocation and embryos were harvested in prewarmed PBS containing calcium chloride and magnesium chloride (D1283 Millipore Sigma). Gross dissection of the embryonic brain rostral to the optic cup (excluding the hindbrain) performed using a Leica stereomicroscope at 12.5× magnification. Tissue pieces were gently dissociated with Accutase and mechanical trituration with a P1000 micropipette. Enzymatic dissociation was stopped by dilution in N2B27 base medium. Dissociated primary NSPCs in suspension were pelleted by centrifugation and re-suspended in complete NSE growth medium for routine monolayer subculture on poly-D-lysine- and laminin-coated T25 flasks.

### Immunofluorescence Microscopy and Proximity Ligation Assays

Cells grown on coated glass coverslips were fixed with 10% neutral formalin (Millipore Sigma HT501128) for 15 min. Cells were permeabilized and blocked with 10% fetal bovine serum (FBS) and 0.1% Triton-x100 in 1X phosphate buffer saline (PBS) for 60 min at room temperature. Primary antibodies were applied for overnight incubation at 4 °C at a dilution of 1:40. Cells were washed twice with 0.1% Tween-20 in PBS and secondary antibodies were applied at a dilution of 1:1000 for 2 hours. Primary and secondary antibodies were diluted in 0.2% FBS, and 0.15% Tween-20 in PBS. Antibody product identifiers and dilutions used for immunofluorescence are listed in **Table S18**. Nuclei were counterstained with 5 ng/mL 4’,6-diamidino-2-phenylindole (DAPI) for 30 seconds. Following five washes in PBS, coverslips were mounted onto slides with Vectashield (Vector Laboratories H100010) and sealed using transparent nail polish. Epifluorescence imaging was performed on a Zeiss Axio Imager Z2 Upright Microscope (ZEISS), equipped with a X-cite Exacte light source (Excelitas Technologies) and a CoolCube 1 camera (MetaSystems). Raw images were processed in FIJI (Schindelin et al., 2012).

For *in situ* protein detection by proximity-dependent DNA ligation assays (Fredriksson et al. 2002), NSPCs were fixed and processed on coverslips as above. Primary antibodies against phospho-serine 5 carboxy terminal domain (S5P CTD) RNA polymerase II (Abcam ab5131; RRID: AB_449369) and SOX2 (R and D Systems MAB2018; RRID: AB_358009) (MAB2018, R&D Systems), were diluted 1:200 and 1:1000 (respectively) into 0.2% FBS/0.1% Triton X-100 in PBS and incubated on coverslips overnight at 4 °C. Following this, coverslips with NSPCs were washed three times in 0.1% Tween-20 in PBS for 5 minutes each. Duolink probes (mouse plus and rabbit minus) were prepared and applied as described by manufacturer’s instructions for the Duolink® PLA kit Cy5 (Millipore-Sigma DUO92013).

Coverslips with NSPCs were mounted in VECTASHIELD Antifade Mounting Medium with DAPI. Images were collected using a Leica TCS SP8 confocal microscope and 100X magnification objective lens with oil immersion at constant exposure. Individual nuclei were identified using Imaris 7.1 (Oxford Instruments; RRID:SCR_007370). 3D models rendered and PLA focal signals were automatically counted per nucleus. All PLA experiments were carried out on at least three independent cultures of NSPCs and a minimum of 100 individual nuclei were imaged per genotype. Single antibody and PLA probe hybridization negative controls were also conducted.

### Intracellular Flow Cytometry

For intracellular antigen staining flow cytometry, cells were incubated with 1 µL of LIVE/DEAD Fixable Dead Cell Stain (Thermo Fisher L34963) per mL of culture media at 37°C for 30 minutes, followed by two washes with PBS. Cultures were dissociated using Accutase, collected, centrifuged at 400g for 4 minutes, and resuspended in >200 µL of 4% PFA for 10 minutes at room temperature. After washing out PFA with PBS, cells were resuspended in 500 µL 0.5% BSA in PBS. A permeabilization and staining buffer (0.5% BSA and 0.1% Triton X-100 in PBS) was prepared. Cells were counted, aliquoted at 0.5 × 10^6^ cells per tube and resuspended in permeabilization and staining buffer for 10 minutes.

Directly conjugated antibodies targeting SOX2 (eFluor570) or an isotype control were added (**Table S18**), and cells were incubated for 1.5 hours at room temperature, protected from light. Cells were washed and resuspended in PBS + 0.5% BSA for data acquisition a BD Fortessa analyzer. Population gating for single live cells and quantification of median SOX2 staining intensity was performed using FlowJo v10 (RRID:SCR_008520).

### RNA Extraction and Reverse-Transcription Quantitative PCR

Total RNA from cultured cells was isolated using RNeasy RNA kit (Qiagen) and processed with TURBO DNA-free kit (Thermo Fisher AM1907). DNase-treated RNA was reverse transcribed with random primers using the high-capacity cDNA synthesis kit (Thermo Fisher 4368814). Target gene expression was monitored by quantitative PCR and normalized to *Gapdh* and *Eef2a* levels with gene-specific primers (oligonucleotide sequences provided in **Table S19**). We monitored expression of *Sox2* by allele-specific reverse transcription quantitative real-time PCR (RT-qPCR) due to the presence of discriminatory SNPs within the *Sox2* transcript, as previously described (Zhou et al. 2014). For each biological replicate, qPCR reactions were performed in technical duplicate using SYBR Select Master mix (Thermo Fisher 4472908) and the CFX 384 Real-Time Detection system (BIO-RAD; RRID:SCR_018057). Expression levels were interpolated from a standard curve dilution series of F1 ESC genomic DNA using CFX Maestro Software. All RNA samples were confirmed not to have DNA contamination because no amplification was observed in reverse transcriptase negative samples.

### Reporter Plasmid Construction

SRR2 and SRR107 were amplified from a Sox2 BAC (RP23-274P9) by PCR using Phusion polymerase (New England Biolabs E0553S) and cloned into the pJET1.2/blunt cloning vector (Thermo Fisher K1232) by Zhou et al. (Zhou et al. 2014). The remaining *Sox2* neural enhancer candidates were amplified from genomic DNA isolated from Bl6 mouse tissue (Jackson Labs) using the DNeasy Blood and Tissue Kit (Qiagen 69504). The 1.8-kb *Sox2* promoter (pSox2) was amplified from mouse genomic DNA by PCR using DreamTaq polymerase (Thermo Fisher EP0702) and ligated into the BanII site to replace the TATA-box minimal promoter of the pGL4.23 vector. All primers used in plasmid construction are provided in **Table S17**. Enhancer sequences were amplified using primers containing 15 bp 5’overhangs homologous to the sequences flanking the NotI restriction site in the pGL4.23 luciferase reporter vector. Amplicons were cloned into the NotI site of the pGL4.23 vector with pSox2 by In-Fusion® cloning (Takara Bio 639650). All plasmids were purified from bacterial culture using Presto™ Midi Plasmid Kit (Geneaid Biotech Inc. PIE025).

### Dual Luciferase Reporter Assay

To evaluate enhancer activity in a luciferase reporter assay, F1 ESCs were seeded on gelatin-coated 96-well plates at a density of 10,000 cells per well. Alternatively, F1 NSPCs were seeded onto PDL/laminin-coated 96-well plates at a density of 50,000 cells per well. After 24 hours, the cells were co-transfected using Lipofectamine 3000 (Thermo Fisher L3000008) with pSox2-pGL4.23 reporter vectors and pGL4.75 encoding Renilla luciferase (Promega E8411 & E6931) at a 50:1 molar ratio. The Renilla plasmid served as an internal control for transfection efficiency in each well. After 24 hours, spent growth medium was replaced with fresh growth medium, according to the cell type transfected. Luciferase activity in cell lysates was assayed 48 hours post-transfection using the Dual Luciferase Reporter Assay kit (Promega E1910), and the Fluoroskan Ascent FL microplate reader (Thermo Fisher Scientific). After background signal correction, the ratio of firefly/Renilla luciferase activity was calculated for each tested enhancer candidate and was normalized to that of the empty vector.

### Chromatin Immunoprecipitation with Sequencing (ChIP-seq) Retrieval and Data Processing

Raw H3K27ac ChIP-seq data from embryonic mouse forebrain tissue samples at E14.5 were obtained from the ENCODE project consortium (ENCODE Project Consortium 2012) with data identifiers provided in **Table S1**. Raw single-end ChIP-seq reads retrieved from the Sequence Read Archive (https://www.ncbi.nlm.nih.gov/sra) or the European Nucleotide Archive (https://www.ebi.ac.uk/ena/browser/home) are listed in Supplemental **Tables S2**, which include data from several studies (Lodato et al. 2013; Quevedo et al. 2019; Phillips-Cremins et al. 2013; Bonev et al. 2017; Bertolini et al. 2019; Pagin et al. 2021; Webb et al. 2013; Estarás et al. 2012; Lodato et al. 2014; Mateo et al. 2015; Sun et al. 2015; Sessa et al. 2017; Braccioli et al. 2018). ChIP-seq reads were assessed for quality, adapter content and duplication rates with FastQC (Andrews 2010), trimmed with Fastp, and aligned to mm10 (Frankish et al. 2019) with Bowtie2 (Langmead and Salzberg 2012), with the following parameters: -N 1 --sensitive- --no-unal --threads 8 -x 700. Removal of duplicated reads and reads mapping to ENCODE mm10 blacklist regions were performed using Samtools (Li et al. 2009). For visualization purposes, we used deepTools bamCoverage (Ramírez et al. 2014) to generate coverage profiles normalized to a depth of one million reads. Peaks were called for each DNA binding protein sample separately using MACS2 narrow peak calling (Zhang et al. 2008).

### RNA-sequencing Library Preparation and Data Processing

Total RNA from cultured NSPCs was isolated and processed in the same way as samples prepared for reverse-transcription quantitative PCR assays. Total RNA was sent to The Centre for Applied Genomics (The Hospital for Sick Children, Toronto) for paired-end rRNA-depleted total RNA-seq (Illumina 2500, 125 bp). Raw RNA-seq reads from the public domain were retrieved from the GEO, which include data from the following studies (Bertolini et al. 2019; Lattke et al. 2021). Raw reads were assessed for quality, adapter content and duplication rates with FastQC and trimmed with Fastp v0.20.0 (S. Chen et al. 2018). Trimmed reads were aligned to mm10 (GENCODE M25) with with STAR v2.9.0a (Dobin et al. 2013), with the following: parameters --alignEndsType Local -- -- outFilterMultimapNmax 20. Exon-mapped reads were quantified using featureCounts (Liao et al. 2014) and imported into R v4.3.2 for differential expression analysis with DESeq2 (Love et al. 2014). Genes with an absolute log2 fold-change > 1 and false discovery rate-adjusted P < 0.05 in total exon-mapped read counts were considered differentially expressed.

For allele-specific RNA-seq, a hybrid version of mm10 was created with SNPsplit v0.3.4 (Krueger and Andrews 2016) using CAST/EiJ SNPs to N-mask discriminatory variants on a 129S1/SvImJ SNP-substituted reference based on GRCh38 from the Mouse Genome Project (dbSNP142). Trimmed reads were aligned to the hybrid N-masked assembly with STAR v2.9.0a, with the following parameters: -- alignEndsType EndToEnd --outSAMattributes NH HI NM MD --outMultimapperOrder Random -- outSAMmultNmax 1. Allele-specific sorting of alignment files was performed using SNPsplit with default parameters. Genes with an absolute log2 fold-change > 0.5 and false discovery rate-adjusted P < 0.05 in allele-sorted read counts using DESeq2 were considered to have significant allelic imbalances. The alignment files of all replicates per sample group were combined using Samtools merge and used to create bedgraphs that were normalized to counts per million mapped reads (CPM) using Deeptools bamCoverage (Ramírez et al. 2014).

### Assay for Transposase Accessible Chromatin with Sequencing (ATAC-seq) Library Preparation and Data Processing

NSPC ATAC-seq multiplex library preparation and next-generation sequencing were performed at the Princess Margaret Genomics Centre, Toronto, Canada (www.pmgenomics.ca). 30,000 – 50,000 Sox2-mCherry^129^ and ΔSRR2-18^129/Cast^ NSPCs were dissociated with Accutase and washed once with 0.04% BSA in PBS. Intact nuclei were isolated and processed using standard ATAC-seq procedures developed by Buenrostro et al. (Buenrostro et al. 2013) with modifications to the lysis and transposition steps based on the OMNI-ATAC protocol by Corces et al. (Corces et al. 2017). Demultiplexed raw ATAC-seq reads were processed using the ENCODE ATAC-seq pipeline (v.1.10.0) established by the Kundaje Lab (RRID:SCR_023100). In this pipeline, MACS2 was used for peak-calling and conserved peaks in biological replicates were filtered using Irreproducible Discovery Rate (IDR; RRID:SCR_017237) (Li et al. 2011). Accessible chromatin regions identified by conservative thresholded IDR peaks were retained for downstream analyses. The alignment files of all replicates per sample group were combined using Samtools merge and used to create bedGraphs that were normalized to counts per million mapped reads (CPM) using Deeptools bamCoverage. IDR peaks were imported into R and annotated to the nearest transcriptional start site of genes using the ChIPpeakanno package (Zhu et al. 2010). Normalization of read counts and analysis of differential accessibility for the union peak regions were performed using DiffBind package (Stark and Brown 2011). Peak intervals with *Q* ≤ 0.00001 and log2 fold change ≥ 1.5 were considered differentially accessible.

### Transcription Factor Motif Enrichment

Transcription factor digital footprint analysis was performed using TOBIAS with standard settings (Bentsen et al. 2020) and a reference compendium of non-redundant mammalian TF motifs from the Jaspar 2022 release (Castro-Mondragon et al. 2022). Non-redundant motifs with an absolute log2 fold-change > 0.2 and *Q* value < 0.01 were considered significantly enriched in each condition. Independent cell culture replicates (n = 3) were merged into a single BAM file for each treatment for Tn5 bias correction and footprint scoring. A consensus peak set was exported from DiffBind. ENCODE mm10 blacklist regions (ENCFF543DDX) were excluded (Amemiya et al. 2019).

### Gene Set Enrichment Analyses

Gene set enrichment analysis on RNA-seq data was performed by ranking genes according to their log2 fold-change and analyzed using the R package ‘clusterProfiler’ (Yu et al. 2012). The function ‘enrichGO’ was used with the following parameters: OrgDb = org.Mm.eg.db, keyType = ‘SYMBOL’, ont = ‘BP’, pAdjustMethod = ‘BH’, pvalueCutoff = 0.01, qvalueCutoff = 0.05.Gene Ontology analysis was performed on narrowPeak files of accessible genomic regions using the R package ‘chipenrich’ (Welch et al. 2014) and the function polyenrich with the following parameters: genesets = GOBP, method=“polyenrich-weighted”, multiAssign=TRUE.

### Data Visualization

Principle component analysis biplots were plotted using PCAtools R package (Blighe and Lun 2020). MA plots for allelic imbalances in gene expression were plotted using DESeq2 (Love et al. 2014). ATAC-seq signal enrichment in accessible chromatin was performed using NGSplot (Shen et al. 2014). Differential gene expression and differential motif scores were plotted using the EnhancedVolcano R package (Blighe et al. 2019). Correlation and clustering heatmaps were plotted using the pheatmap R package (Kolde 2012).

### Statistical Analysis

Data were analyzed with R version 4.2.3 (R Core Team 2020) and the Tidyverse package (Wickham et al. 2019). Statistical methods, *P* values or adjusted *P* values for each comparison are listed in the figure legend and/or in the corresponding results section. Assumptions of normality and homogeneity of variance were assessed with Shapiro-Wilk and Fligner-Killeen tests. For all experiments, sample size was determined empirically. Investigators were not blinded to allocation during experiments or outcome assessments.

### Data and Resource Availability

All cell lines and plasmids described are available upon request. The authors affirm that all data necessary for confirming the conclusions of this article are represented fully within the article and its tables and figures, supporting information, and the following data repositories. Raw sequencing data and processed bedGraph or narrowPeak files from this work were submitted to the GEO repository under the reference series GSE237778. RT-qPCR, luciferase reporter, PLA immunofluorescence, flow cytometry data are deposited in Zenodo (DOI: 10.5281/zenodo.12750519). This work does not use any original code.

## Competing Interests Statement

The authors declare no competing interests.

## Acknowledgments

The authors thank members of the Mitchell group for helpful discussions. This work was supported by the Canadian Institutes of Health Research (FRN PJT153186 and PJT180312), the Canada Foundation for Innovation, and the Ontario Ministry of Research and Innovation (operating and infrastructure grants held by J.A.M.). Salary support for S. D. M. was provided by Canadian Institutes of Health Research Project Grant (FRN 153186). Salary support for I.C.T was provided by a fellowship from the University of Toronto. Salary support for Z. E. G. was provided by a Canadian Institutes of Health Research fellowship. Studentship funding was provided by the Natural Science and Engineering Research Council of Canada (CGS Doctoral held by V. M. S.) This research was enabled in part by support provided by Compute Ontario (https://www.computeontario.ca/) and the Digital Research Alliance of Canada (https://www.alliancecan.ca). We thank the ENCODE Consortium for data sets queried here. We also thank Alan M. Moses for allowing A.G.D. to use computational resources while in his lab.

## Author Contributions

I.C.T., S.D.M., and J.A.M. conceived the project and designed the experiments. I.C.T, S.D.M., V.M.S., L.L., Z.E.G., N.G., M.C., and R.B.D. made experimental contributions to the gene expression analysis, generation of deleted ESC lines, NSPC derivation and maintenance, plasmid construction and reporter assays, sample preparation for fluorescence microscopy, confocal imaging, RNA-seq and ATAC-seq data processing. I.C.T and A.G.D curated and formally analyzed all experimental data. I.C.T. generated data visualizations and wrote the original draft. J.A.M. acquired funding and provided resources, administration, and supervision to this project. Review and editing of the manuscript were carried out by all co-authors.

## Notes

### Competing Interest Statement

The authors have declared no competing interest.

### Summary of Updates

Characterization of the NSC line obtained by differentiating mouse ESCs was added to Figure 3 as panel D. Luciferase data was added for NSPCs isolated from the developing mouse brain in Figure 2C. Sox2 protein levels were added for enhancer deleted lines in Figure 4B.

## References

Alder O, Cullum R, Lee S, Kan AC, Wei W, Yi Y, Garside VC, Bilenky M, Griffith M, Morrissy AS. 2014. Hippo signaling influences HNF4A and FOXA2 enhancer switching during hepatocyte differentiation. Cell Rep. 9(1):261–271.

Amador-Arjona A, Cimadamore F, Huang C-T, Wright R, Lewis S, Gage FH, Terskikh AV. 2015. SOX2 primes the epigenetic landscape in neural precursors enabling proper gene activation during hippocampal neurogenesis. Proc Natl Acad Sci. 112(15). 10.1073/pnas.1421480112

Amemiya HM, Kundaje A, Boyle AP. 2019. The ENCODE Blacklist: Identification of Problematic Regions of the Genome. Sci Rep. 9(1):9354. 10.1038/s41598-019-45839-z

Andreu-Agullo C, Maurin T, Thompson CB, Lai EC. 2012. Ars2 maintains neural stem-cell identity through direct transcriptional activation of Sox2. Nature. 481(7380):195–198. 10.1038/nature10712

Andrews S. 2010. FastQC: a quality control tool for high throughput sequence data.

Avilion AA, Nicolis SK, Pevny LH, Perez L, Vivian N, Lovell-Badge R. 2003. Multipotent cell lineages in early mouse development depend on SOX2 function. Genes Dev. 17(1):126–140. 10.1101/gad.224503

Bailey PJ, Klos JM, Andersson E, Karlén M, Källström M, Ponjavic J, Muhr J, Lenhard B, Sandelin A, Ericson J. 2006. A global genomic transcriptional code associated with CNS-expressed genes. Exp Cell Res. 312(16):3108–3119. 10.1016/j.yexcr.2006.06.017

Banerji J, Rusconi S, Schaffner W. 1981. Expression of a β-globin gene is enhanced by remote SV40 DNA sequences. Cell. 27(2, Part 1):299–308. 10.1016/0092-8674(81)90413-X

Baumann V, Wiesbeck M, Breunig CT, Braun JM, Köferle A, Ninkovic J, Götz M, Stricker SH. 2019. Targeted removal of epigenetic barriers during transcriptional reprogramming. Nat Commun. 10(1):2119. 10.1038/s41467-019-10146-8

Beagan JA, Duong MT, Titus KR, Zhou L, Cao Z, Ma J, Lachanski CV, Gillis DR, Phillips-Cremins JE. 2017. YY1 and CTCF orchestrate a 3D chromatin looping switch during early neural lineage commitment. Genome Res. 27(7):1139–1152. 10.1101/gr.215160.116

Beagan JA, Gilgenast TG, Kim J, Plona Z, Norton HK, Hu G, Hsu SC, Shields EJ, Lyu X, Apostolou E, et al. 2016. Local Genome Topology Can Exhibit an Incompletely Rewired 3D-Folding State during Somatic Cell Reprogramming. Cell Stem Cell. 18(5):611–624. 10.1016/j.stem.2016.04.004

Bejjani F, Evanno E, Zibara K, Piechaczyk M, Jariel-Encontre I. 2019. The AP-1 transcriptional complex: Local switch or remote command? Biochim Biophys Acta BBA - Rev Cancer. 1872(1):11–23. 10.1016/j.bbcan.2019.04.003

Bentsen M, Goymann P, Schultheis H, Klee K, Petrova A, Wiegandt R, Fust A, Preussner J, Kuenne C, Braun T. 2020. ATAC-seq footprinting unravels kinetics of transcription factor binding during zygotic genome activation. Nat Commun. 11(1):4267.

Bertacchi M, Pandolfini L, Murenu E, Viegi A, Capsoni S, Cellerino A, Messina A, Casarosa S, Cremisi F. 2013. The positional identity of mouse ES cell-generated neurons is affected by BMP signaling. Cell Mol Life Sci CMLS. 70(6):1095–1111. 10.1007/s00018-012-1182-3

Bertolini JA, Favaro R, Zhu Y, Pagin M, Ngan CY, Wong CH, Tjong H, Vermunt MW, Martynoga B, Barone C, et al. 2019. Mapping the Global Chromatin Connectivity Network for Sox2 Function in Neural Stem Cell Maintenance. Cell Stem Cell. 24(3):462–476.e6. 10.1016/j.stem.2019.02.004

Blassberg R, Patel H, Watson T, Gouti M, Metzis V, Delás MJ, Briscoe J. 2022. Sox2 levels regulate the chromatin occupancy of WNT mediators in epiblast progenitors responsible for vertebrate body formation. Nat Cell Biol. 24(5):633–644. 10.1038/s41556-022-00910-2

Blighe K, Lun A. 2020. PCAtools: everything principal components analysis. R Package Version. 2(0).

Blighe K, Rana S, Lewis M. 2019. EnhancedVolcano: Publication-ready volcano plots with enhanced colouring and labeling. R Package Version. 1(0).

Bonev B, Mendelson Cohen N, Szabo Q, Fritsch L, Papadopoulos GL, Lubling Y, Xu X, Lv X, Hugnot J- P, Tanay A, Cavalli G. 2017. Multiscale 3D Genome Rewiring during Mouse Neural Development. Cell. 171(3):557–572.e24. 10.1016/j.cell.2017.09.043

Braccioli L, Vervoort SJ, Puma G, Nijboer CH, Coffer PJ. 2018. SOX4 inhibits oligodendrocyte differentiation of embryonic neural stem cells in vitro by inducing Hes5 expression. Stem Cell Res. 33:110–119. 10.1016/j.scr.2018.10.005

Brandenberger R, Wei H, Zhang S, Lei S, Murage J, Fisk GJ, Li Y, Xu C, Fang R, Guegler K, et al. 2004. Transcriptome characterization elucidates signaling networks that control human ES cell growth and differentiation. Nat Biotechnol. 22(6):707–716. 10.1038/nbt971

Brosh R, Coelho C, Ribeiro-dos-Santos AM, Ellis G, Hogan MS, Ashe HJ, Somogyi N, Ordoñez R, Luther RD, Huang E. 2023. Synthetic regulatory genomics uncovers enhancer context dependence at the Sox2 locus. Mol Cell. 83(7):1140–1152.

Buenrostro JD, Giresi PG, Zaba LC, Chang HY, Greenleaf WJ. 2013. Transposition of native chromatin for fast and sensitive epigenomic profiling of open chromatin, DNA-binding proteins and nucleosome position. Nat Methods. 10(12):1213–1218. 10.1038/nmeth.2688

Carter D, Chakalova L, Osborne CS, Dai Y, Fraser P. 2002. Long-range chromatin regulatory interactions in vivo. Nat Genet. 32(4):623–626.

Castro-Mondragon JA, Riudavets-Puig R, Rauluseviciute I, Berhanu Lemma R, Turchi L, Blanc-Mathieu R, Lucas J, Boddie P, Khan A, Manosalva Pérez N. 2022. JASPAR 2022: the 9th release of the open-access database of transcription factor binding profiles. Nucleic Acids Res. 50(D1):D165–D173.

Catena R, Tiveron C, Ronchi A, Porta S, Ferri A, Tatangelo L, Cavallaro M, Favaro R, Ottolenghi S, Reinbold R, et al. 2004. Conserved POU Binding DNA Sites in the Sox2 Upstream Enhancer Regulate Gene Expression in Embryonic and Neural Stem Cells. J Biol Chem. 279(40):41846–41857. 10.1074/jbc.M405514200

Cavallaro M, Mariani J, Lancini C, Latorre E, Caccia R, Gullo F, Valotta M, DeBiasi S, Spinardi L, Ronchi A, et al. 2008. Impaired generation of mature neurons by neural stem cells from hypomorphic Sox2 mutants. Development. 135(3):541–557. 10.1242/dev.010801

Chakraborty S, Kopitchinski N, Zuo Z, Eraso A, Awasthi P, Chari R, Mitra A, Tobias IC, Moorthy SD, Dale RK, et al. 2023. Enhancer–promoter interactions can bypass CTCF-mediated boundaries and contribute to phenotypic robustness. Nat Genet. 55(2):280–290. 10.1038/s41588-022-01295-6

Chawengsaksophak K, de Graaff W, Rossant J, Deschamps J, Beck F. 2004. Cdx2 is essential for axial elongation in mouse development. Proc Natl Acad Sci U S A. 101(20):7641–7645. 10.1073/pnas.0401654101

Chen AF, Liu AJ, Krishnakumar R, Freimer JW, DeVeale B, Blelloch R. 2018. GRHL2-Dependent Enhancer Switching Maintains a Pluripotent Stem Cell Transcriptional Subnetwork after Exit from Naive Pluripotency. Cell Stem Cell. 23(2):226–238.e4. 10.1016/j.stem.2018.06.005

Chen C, Morris Q, Mitchell JA. 2012. Enhancer identification in mouse embryonic stem cells using integrative modeling of chromatin and genomic features. BMC Genomics. 13(1):152. 10.1186/1471-2164-13-152

Chen S, Zhou Y, Chen Y, Gu J. 2018. fastp: an ultra-fast all-in-one FASTQ preprocessor. Bioinformatics. 34(17):i884–i890. 10.1093/bioinformatics/bty560

Chen X, Xu H, Yuan P, Fang F, Huss M, Vega VB, Wong E, Orlov YL, Zhang W, Jiang J, et al. 2008. Integration of External Signaling Pathways with the Core Transcriptional Network in Embryonic Stem Cells. Cell. 133(6):1106–1117. 10.1016/j.cell.2008.04.043

Chi H, Sarkisian MR, Rakic P, Flavell RA. 2005. Loss of mitogen-activated protein kinase kinase kinase 4 (MEKK4) results in enhanced apoptosis and defective neural tube development. Proc Natl Acad Sci U S A. 102(10):3846–3851. 10.1073/pnas.0500026102

Corces MR, Trevino AE, Hamilton EG, Greenside PG, Sinnott-Armstrong NA, Vesuna S, Satpathy AT, Rubin AJ, Montine KS, Wu B. 2017. An improved ATAC-seq protocol reduces background and enables interrogation of frozen tissues. Nat Methods. 14(10):959–962.

Creyghton MP, Cheng AW, Welstead GG, Kooistra T, Carey BW, Steine EJ, Hanna J, Lodato MA, Frampton GM, Sharp PA, et al. 2010. Histone H3K27ac separates active from poised enhancers and predicts developmental state. Proc Natl Acad Sci. 107(50):21931–21936. 10.1073/pnas.1016071107

Delás MJ, Kalaitzis CM, Fawzi T, Demuth M, Zhang I, Stuart HT, Costantini E, Ivanovitch K, Tanaka EM, Briscoe J. 2023. Developmental cell fate choice in neural tube progenitors employs two distinct cis-regulatory strategies. Dev Cell. 58(1):3–17.e8. 10.1016/j.devcel.2022.11.016

Dhaliwal NK, Abatti LE, Mitchell JA. 2019. KLF4 protein stability regulated by interaction with pluripotency transcription factors overrides transcriptional control. Genes Dev. 33(15–16):1069–1082. 10.1101/gad.324319.119

Dhaliwal NK, Miri K, Davidson S, Tamim El Jarkass H, Mitchell JA. 2018. KLF4 Nuclear Export Requires ERK Activation and Initiates Exit from Naive Pluripotency. Stem Cell Rep. 10(4):1308–1323. 10.1016/j.stemcr.2018.02.007

Diez Del Corral R, Morales AV. 2017. The Multiple Roles of FGF Signaling in the Developing Spinal Cord. Front Cell Dev Biol. 5:58. 10.3389/fcell.2017.00058

Ding Q, Lee Y-K, Schaefer EA, Peters DT, Veres A, Kim K, Kuperwasser N, Motola DL, Meissner TB, Hendriks WT. 2013. A TALEN genome-editing system for generating human stem cell-based disease models. Cell Stem Cell. 12(2):238–251.

Dobin A, Davis CA, Schlesinger F, Drenkow J, Zaleski C, Jha S, Batut P, Chaisson M, Gingeras TR. 2013. STAR: ultrafast universal RNA-seq aligner. Bioinformatics. 29(1):15–21. 10.1093/bioinformatics/bts635

Dodonova SO, Zhu F, Dienemann C, Taipale J, Cramer P. 2020. Nucleosome-bound SOX2 and SOX11 structures elucidate pioneer factor function. Nature. 580(7805):669–672.

ENCODE Project Consortium. 2012. An integrated encyclopedia of DNA elements in the human genome. Nature. 489(7414):57.

Ensini M, Tsuchida TN, Belting HG, Jessell TM. 1998. The control of rostrocaudal pattern in the developing spinal cord: specification of motor neuron subtype identity is initiated by signals from paraxial mesoderm. Dev Camb Engl. 125(6):969–982. 10.1242/dev.125.6.969

Estarás C, Akizu N, García A, Beltrán S, de la Cruz X, Martínez-Balbás MA. 2012. Genome-wide analysis reveals that Smad3 and JMJD3 HDM co-activate the neural developmental program. Dev Camb Engl. 139(15):2681–2691. 10.1242/dev.078345

Favaro R, Valotta M, Ferri ALM, Latorre E, Mariani J, Giachino C, Lancini C, Tosetti V, Ottolenghi S, Taylor V, Nicolis SK. 2009. Hippocampal development and neural stem cell maintenance require Sox2-dependent regulation of Shh. Nat Neurosci. 12(10):1248–1256. 10.1038/nn.2397

Ferri ALM, Cavallaro M, Braida D, Di Cristofano A, Canta A, Vezzani A, Ottolenghi S, Pandolfi PP, Sala M, DeBiasi S, Nicolis SK. 2004. Sox2 deficiency causes neurodegeneration and impaired neurogenesis in the adult mouse brain. Development. 131(15):3805–3819. 10.1242/dev.01204

Foshay KM, Gallicano GI. 2008. Regulation of Sox2 by STAT3 initiates commitment to the neural precursor cell fate. Stem Cells Dev. 17(2):269–278.

Frankish A, Diekhans M, Ferreira A-M, Johnson R, Jungreis I, Loveland J, Mudge JM, Sisu C, Wright J, Armstrong J, et al. 2019. GENCODE reference annotation for the human and mouse genomes. Nucleic Acids Res. 47(D1):D766–D773. 10.1093/nar/gky955

Fredriksson S, Gullberg M, Jarvius J, Olsson C, Pietras K, Gústafsdóttir SM, Östman A, Landegren U. 2002. Protein detection using proximity-dependent DNA ligation assays. Nat Biotechnol. 20(5):473– 477.

Garcia ADR, Doan NB, Imura T, Bush TG, Sofroniew MV. 2004. GFAP-expressing progenitors are the principal source of constitutive neurogenesis in adult mouse forebrain. Nat Neurosci. 7(11):1233–1241. 10.1038/nn1340

Götz M, Barde Y-A. 2005. Radial glial cells defined and major intermediates between embryonic stem cells and CNS neurons. Neuron. 46(3):369–372. 10.1016/j.neuron.2005.04.012

Graham V, Khudyakov J, Ellis P, Pevny L. 2003. SOX2 functions to maintain neural progenitor identity. Neuron. 39(5):749–765.

Gubbay J, Collignon J, Koopman P, Capel B, Economou A, Münsterberg A, Vivian N, Goodfellow P, Lovell-Badge R. 1990. A gene mapping to the sex-determining region of the mouse Y chromosome is a member of a novel family of embryonically expressed genes. Nature. 346(6281):245–250. 10.1038/346245a0

Hagey DW, Klum S, Kurtsdotter I, Zaouter C, Topcic D, Andersson O, Bergsland M, Muhr J. 2018. SOX2 regulates common and specific stem cell features in the CNS and endoderm derived organs. PLoS Genet. 14(2):e1007224. 10.1371/journal.pgen.1007224

Hagey DW, Zaouter C, Combeau G, Lendahl MA, Andersson O, Huss M, Muhr J. 2016. Distinct transcription factor complexes act on a permissive chromatin landscape to establish regionalized gene expression in CNS stem cells. Genome Res. 26(7):908–917. 10.1101/gr.203513.115

Hallmann A-L, Araúzo-Bravo MJ, Zerfass C, Senner V, Ehrlich M, Psathaki OE, Han DW, Tapia N, Zaehres H, Schöler HR, et al. 2016. Comparative transcriptome analysis in induced neural stem cells reveals defined neural cell identities in vitro and after transplantation into the adult rodent brain. Stem Cell Res. 16(3):776–781. 10.1016/j.scr.2016.04.015

Hatada I, Namihira M, Morita S, Kimura M, Horii T, Nakashima K. 2008. Astrocyte-Specific Genes Are Generally Demethylated in Neural Precursor Cells Prior to Astrocytic Differentiation. PLOS ONE. 3(9):e3189. 10.1371/journal.pone.0003189

Hay D, Hughes JR, Babbs C, Davies JOJ, Graham BJ, Hanssen LLP, Kassouf MT, Oudelaar AM, Sharpe JA, Suciu MC, et al. 2016. Genetic dissection of the α-globin super-enhancer in vivo. Nat Genet. 48(8):895–903. 10.1038/ng.3605

Heintzman ND, Hon GC, Hawkins RD, Kheradpour P, Stark A, Harp LF, Ye Z, Lee LK, Stuart RK, Ching CW, et al. 2009. Histone modifications at human enhancers reflect global cell-type-specific gene expression. Nature. 459(7243):108–112. 10.1038/nature07829

Hsieh T-HS, Cattoglio C, Slobodyanyuk E, Hansen AS, Darzacq X, Tjian R. 2022. Enhancer–promoter interactions and transcription are largely maintained upon acute loss of CTCF, cohesin, WAPL or YY1. Nat Genet. 54(12):1919–1932. 10.1038/s41588-022-01223-8

Huang J, Liu X, Li D, Shao Z, Cao H, Zhang Y, Trompouki E, Bowman TV, Zon LI, Yuan G-C, et al. 2016. Dynamic Control of Enhancer Repertoires Drives Lineage and Stage-Specific Transcription during Hematopoiesis. Dev Cell. 36(1):9–23. 10.1016/j.devcel.2015.12.014

Ivanova N, Dobrin R, Lu R, Kotenko I, Levorse J, DeCoste C, Schafer X, Lun Y, Lemischka IR. 2006. Dissecting self-renewal in stem cells with RNA interference. Nature. 442(7102):533–538. 10.1038/nature04915

Iwafuchi-Doi M, Donahue G, Kakumanu A, Watts JA, Mahony S, Pugh BF, Lee D, Kaestner KH, Zaret KS. 2016. The Pioneer Transcription Factor FoxA Maintains an Accessible Nucleosome Configuration at Enhancers for Tissue-Specific Gene Activation. Mol Cell. 62(1):79–91. 10.1016/j.molcel.2016.03.001

Iwafuchi-Doi M, Yoshida Y, Onichtchouk D, Leichsenring M, Driever W, Takemoto T, Uchikawa M, Kamachi Y, Kondoh H. 2011. The Pou5f1/Pou3f-dependent but SoxB-independent regulation of conserved enhancer N2 initiates Sox2 expression during epiblast to neural plate stages in vertebrates. Dev Biol. 352(2):354–366. 10.1016/j.ydbio.2010.12.027

Johe KK, Hazel TG, Muller T, Dugich-Djordjevic MM, McKay RD. 1996. Single factors direct the differentiation of stem cells from the fetal and adult central nervous system. Genes Dev. 10(24):3129– 3140. 10.1101/gad.10.24.3129

Josephson R, Muller T, Pickel J, Okabe S, Reynolds K, Turner PA, Zimmer A, McKay RD. 1998. POU transcription factors control expression of CNS stem cell-specific genes. Development. 125(16):3087– 3100.

Kagey MH, Newman JJ, Bilodeau S, Zhan Y, Orlando DA, van Berkum NL, Ebmeier CC, Goossens J, Rahl PB, Levine SS, et al. 2010. Mediator and cohesin connect gene expression and chromatin architecture. Nature. 467(7314):430–435. 10.1038/nature09380

Kelberman D, Rizzoti K, Avilion A, Bitner-Glindzicz M, Cianfarani S, Collins J, Chong WK, Kirk JMW, Achermann JC, Ross R, et al. 2006. Mutations within Sox2/SOX2 are associated with abnormalities in the hypothalamo-pituitary-gonadal axis in mice and humans. J Clin Invest. 116(9):2442–2455. 10.1172/JCI28658

Kieffer-Kwon K-R, Tang Z, Mathe E, Qian J, Sung M-H, Li G, Resch W, Baek S, Pruett N, Grøntved L, et al. 2013. Interactome Maps of Mouse Gene Regulatory Domains Reveal Basic Principles of Transcriptional Regulation. Cell. 155(7):1507–1520. 10.1016/j.cell.2013.11.039

Kolde R. 2012. Pheatmap: pretty heatmaps. R Package Version. 1(2):726.

Komarnitsky P, Cho E-J, Buratowski S. 2000. Different phosphorylated forms of RNA polymerase II and associated mRNA processing factors during transcription. Genes Dev. 14(19):2452–2460.

Kopp JL, Ormsbee BD, Desler M, Rizzino A. 2008. Small increases in the level of Sox2 trigger the differentiation of mouse embryonic stem cells. Stem Cells. 26(4):903–911.

Krueger F, Andrews SR. 2016. SNPsplit: Allele-specific splitting of alignments between genomes with known SNP genotypes. F1000Res 5, 1479.

Langer L, Taranova O, Sulik K, Pevny L. 2012. SOX2 hypomorphism disrupts development of the prechordal floor and optic cup. Mech Dev. 129(1–4):1–12. 10.1016/j.mod.2012.04.001

Langmead B, Salzberg SL. 2012. Fast gapped-read alignment with Bowtie 2. Nat Methods. 9(4):357–359. 10.1038/nmeth.1923

Lattke M, Goldstone R, Ellis JK, Boeing S, Jurado-Arjona J, Marichal N, MacRae JI, Berninger B, Guillemot F. 2021. Extensive transcriptional and chromatin changes underlie astrocyte maturation in vivo and in culture. Nat Commun. 12(1):4335. 10.1038/s41467-021-24624-5

Lendahl U, Zimmerman LB, McKay RD. 1990. CNS stem cells express a new class of intermediate filament protein. Cell. 60(4):585–595.

Li H, Collado M, Villasante A, Matheu A, Lynch CJ, Cañamero M, Rizzoti K, Carneiro C, Martínez G, Vidal A, et al. 2012. p27Kip1 Directly Represses Sox2 during Embryonic Stem Cell Differentiation. Cell Stem Cell. 11(6):845–852. 10.1016/j.stem.2012.09.014

Li H, Handsaker B, Wysoker A, Fennell T, Ruan J, Homer N, Marth G, Abecasis G, Durbin R, 1000 Genome Project Data Processing Subgroup. 2009. The Sequence Alignment/Map format and SAMtools. Bioinformatics. 25(16):2078–2079. 10.1093/bioinformatics/btp352

Li Q, Brown JB, Huang H, Bickel PJ. 2011. Measuring reproducibility of high-throughput experiments.

Liao Y, Smyth GK, Shi W. 2014. featureCounts: an efficient general purpose program for assigning sequence reads to genomic features. Bioinformatics. 30(7):923–930. 10.1093/bioinformatics/btt656

Lodato MA, Ng CW, Wamstad JA, Cheng AW, Thai KK, Fraenkel E, Jaenisch R, Boyer LA. 2013. SOX2 Co-Occupies Distal Enhancer Elements with Distinct POU Factors in ESCs and NPCs to Specify Cell State. Barsh GS, editor. PLoS Genet. 9(2):e1003288. 10.1371/journal.pgen.1003288

Lodato S, Molyneaux BJ, Zuccaro E, Goff LA, Chen H-H, Yuan W, Meleski A, Takahashi E, Mahony S, Rinn JL, et al. 2014. Gene co-regulation by Fezf2 selects neurotransmitter identity and connectivity of corticospinal neurons. Nat Neurosci. 17(8):1046–1054. 10.1038/nn.3757

Loh Y-H, Wu Q, Chew J-L, Vega VB, Zhang W, Chen X, Bourque G, George J, Leong B, Liu J, et al. 2006. The Oct4 and Nanog transcription network regulates pluripotency in mouse embryonic stem cells. Nat Genet. 38(4):431–440. 10.1038/ng1760

Love M, Anders S, Huber W. 2014. Differential analysis of count data–the DESeq2 package. Genome Biol. 15(550):10–1186.

Mali P, Yang L, Esvelt KM, Aach J, Guell M, DiCarlo JE, Norville JE, Church GM. 2013. RNA-guided human genome engineering via Cas9. Science. 339(6121):823–826.

Marqués-Torrejón MÁ, Porlan E, Banito A, Gómez-Ibarlucea E, Lopez-Contreras AJ, Fernández-Capetillo Ó, Vidal A, Gil J, Torres J, Fariñas I. 2013. Cyclin-Dependent Kinase Inhibitor p21 Controls Adult Neural Stem Cell Expansion by Regulating Sox2 Gene Expression. Cell Stem Cell. 12(1):88–100. 10.1016/j.stem.2012.12.001

Martinez-Ara M, Comoglio F, van Arensbergen J, van Steensel B. 2022. Systematic analysis of intrinsic enhancer-promoter compatibility in the mouse genome. Mol Cell. 82(13):2519–2531.e6. 10.1016/j.molcel.2022.04.009

Masui S, Nakatake Y, Toyooka Y, Shimosato D, Yagi R, Takahashi K, Okochi H, Okuda A, Matoba R, Sharov AA, et al. 2007. Pluripotency governed by Sox2 via regulation of Oct3/4 expression in mouse embryonic stem cells. Nat Cell Biol. 9(6):625–635. 10.1038/ncb1589

Mateo JL, van den Berg DL, Haeussler M, Drechsel D, Gaber ZB, Castro DS, Robson P, Lu QR, Crawford GE, Flicek P. 2015. Characterization of the neural stem cell gene regulatory network identifies OLIG2 as a multifunctional regulator of self-renewal. Genome Res. 25(1):41–56.

Mathis L, François Nicolas J. 2000. Different clonal dispersion in the rostral and caudal mouse central nervous system. Development. 127(6):1277–1290.

Mazzoni EO, Mahony S, Peljto M, Patel T, Thornton SR, McCuine S, Reeder C, Boyer LA, Young RA, Gifford DK, Wichterle H. 2013. Saltatory remodeling of Hox chromatin in response to rostrocaudal patterning signals. Nat Neurosci. 16(9):1191–1198. 10.1038/nn.3490

Metzis V, Steinhauser S, Pakanavicius E, Gouti M, Stamataki D, Ivanovitch K, Watson T, Rayon T, Mousavy Gharavy SN, Lovell-Badge R, et al. 2018. Nervous System Regionalization Entails Axial Allocation before Neural Differentiation. Cell. 175(4):1105–1118.e17. 10.1016/j.cell.2018.09.040

Miyagi S, Masui S, Niwa H, Saito T, Shimazaki T, Okano H, Nishimoto M, Muramatsu M, Iwama A, Okuda A. 2008. Consequence of the loss of Sox2 in the developing brain of the mouse. FEBS Lett. 582(18):2811–2815. 10.1016/j.febslet.2008.07.011

Miyagi S, Nishimoto M, Saito T, Ninomiya M, Sawamoto K, Okano H, Muramatsu M, Oguro H, Iwama A, Okuda A. 2006. The Sox2 Regulatory Region 2 Functions as a Neural Stem Cell-specific Enhancer in the Telencephalon. J Biol Chem. 281(19):13374–13381. 10.1074/jbc.M512669200

Miyagi S, Saito T, Mizutani K, Masuyama N, Gotoh Y, Iwama A, Nakauchi H, Masui S, Niwa H, Nishimoto M, et al. 2004. The Sox-2 Regulatory Regions Display Their Activities in Two Distinct Types of Multipotent Stem Cells. Mol Cell Biol. 24(10):4207–4220. 10.1128/MCB.24.10.4207-4220.2004

Mlynarczyk-Evans S, Royce-Tolland M, Alexander MK, Andersen AA, Kalantry S, Gribnau J, Panning B. 2006. X chromosomes alternate between two states prior to random X-inactivation. PLoS Biol. 4(6):e159.

Molofsky AV, Pardal R, Iwashita T, Park I-K, Clarke MF, Morrison SJ. 2003. Bmi-1 dependence distinguishes neural stem cell self-renewal from progenitor proliferation. Nature. 425(6961):962–967. 10.1038/nature02060

Moncho-Amor V, Chakravarty P, Galichet C, Matheu A, Lovell-Badge R, Rizzoti K. 2021. SOX2 is required independently in both stem and differentiated cells for pituitary tumorigenesis in p27-null mice. Proc Natl Acad Sci. 118(7):e2017115118. 10.1073/pnas.2017115118

Moorthy SD, Davidson S, Shchuka VM, Singh G, Malek-Gilani N, Langroudi L, Martchenko A, So V, Macpherson NN, Mitchell JA. 2017. Enhancers and super-enhancers have an equivalent regulatory role in embryonic stem cells through regulation of single or multiple genes. Genome Res. 27(2):246–258. 10.1101/gr.210930.116

Moorthy SD, Mitchell JA. 2016. Generating CRISPR/Cas9 Mediated Monoallelic Deletions to Study Enhancer Function in Mouse Embryonic Stem Cells. J Vis Exp.(110):53552. 10.3791/53552

Osterwalder M, Barozzi I, Tissières V, Fukuda-Yuzawa Y, Mannion BJ, Afzal SY, Lee EA, Zhu Y, Plajzer-Frick I, Pickle CS, et al. 2018. Enhancer redundancy provides phenotypic robustness in mammalian development. Nature. 554(7691):239–243. 10.1038/nature25461

Pagin M, Pernebrink M, Giubbolini S, Barone C, Sambruni G, Zhu Y, Chiara M, Ottolenghi S, Pavesi G, Wei C-L. 2021. Sox2 controls neural stem cell self-renewal through a Fos-centered gene regulatory network. Stem Cells. 39(8):1107–1119.

Patel JR, McCandless EE, Dorsey D, Klein RS. 2010. CXCR4 promotes differentiation of oligodendrocyte progenitors and remyelination. Proc Natl Acad Sci. 107(24):11062–11067. 10.1073/pnas.1006301107

Phillips-Cremins JE, Sauria MEG, Sanyal A, Gerasimova TI, Lajoie BR, Bell JSK, Ong C-T, Hookway TA, Guo C, Sun Y, et al. 2013. Architectural Protein Subclasses Shape 3D Organization of Genomes during Lineage Commitment. Cell. 153(6):1281–1295. 10.1016/j.cell.2013.04.053

Platania A, Erb C, Barbieri M, Molcrette B, Grandgirard E, de Kort MA, Meaburn K, Taylor T, Shchuka VM, Kocanova S. 2023. Competition between transcription and loop extrusion modulates promoter and enhancer dynamics. BioRxiv.:2023–04.

Que J, Okubo T, Goldenring JR, Nam K-T, Kurotani R, Morrisey EE, Taranova O, Pevny LH, Hogan BLM. 2007. Multiple dose-dependent roles for Sox2 in the patterning and differentiation of anterior foregut endoderm. Development. 134(13):2521–2531. 10.1242/dev.003855

Quevedo M, Meert L, Dekker MR, Dekkers DHW, Brandsma JH, van den Berg DLC, Ozgür Z, van IJcken WFJ, Demmers J, Fornerod M, Poot RA. 2019. Mediator complex interaction partners organize the transcriptional network that defines neural stem cells. Nat Commun. 10(1):2669. 10.1038/s41467-019-10502-8

R Core Team. 2020. R Core Team R: a language and environment for statistical computing. Found Stat Comput.

Rada-Iglesias A, Bajpai R, Swigut T, Brugmann SA, Flynn RA, Wysocka J. 2011. A unique chromatin signature uncovers early developmental enhancers in humans. Nature. 470(7333):279–283. 10.1038/nature09692

Ramírez F, Dündar F, Diehl S, Grüning BA, Manke T. 2014. deepTools: a flexible platform for exploring deep-sequencing data. Nucleic Acids Res. 42(W1):W187–W191. 10.1093/nar/gku365

Reynolds BA, Weiss S. 1992. Generation of Neurons and Astrocytes from Isolated Cells of the Adult Mammalian Central Nervous System. Science. 255(5052):1707–1710. 10.1126/science.1553558

Rhodes CT, Thompson JJ, Mitra A, Asokumar D, Lee DR, Lee DJ, Zhang Y, Jason E, Dale RK, Rocha PP, Petros TJ. 2022. An epigenome atlas of neural progenitors within the embryonic mouse forebrain. Nat Commun. 13(1):4196. 10.1038/s41467-022-31793-4

Ribes V, Le Roux I, Rhinn M, Schuhbaur B, Dollé P. 2009. Early mouse caudal development relies on crosstalk between retinoic acid,Shh and Fgf signalling pathways. Development. 136(4):665–676. 10.1242/dev.016204

Roadmap Epigenomics Consortium, Kundaje A, Meuleman W, Ernst J, Bilenky M, Yen A, Heravi-Moussavi A, Kheradpour P, Zhang Z, Wang J, et al. 2015. Integrative analysis of 111 reference human epigenomes. Nature. 518(7539):317–330. 10.1038/nature14248

Rosenberg AB, Roco CM, Muscat RA, Kuchina A, Sample P, Yao Z, Graybuck LT, Peeler DJ, Mukherjee S, Chen W, et al. 2018. Single-cell profiling of the developing mouse brain and spinal cord with split-pool barcoding. Science. 360(6385):176–182. 10.1126/science.aam8999

Sabapathy K, Jochum W, Hochedlinger K, Chang L, Karin M, Wagner EF. 1999. Defective neural tube morphogenesis and altered apoptosis in the absence of both JNK1 and JNK2. Mech Dev. 89(1–2):115–124. 10.1016/s0925-4773(99)00213-0

Sandberg M, Källström M, Muhr J. 2005. Sox21 promotes the progression of vertebrate neurogenesis. Nat Neurosci. 8(8):995–1001. 10.1038/nn1493

Sanyal A, Lajoie BR, Jain G, Dekker J. 2012. The long-range interaction landscape of gene promoters. Nature. 489(7414):109–113. 10.1038/nature11279

Schoenfelder S, Furlan-Magaril M, Mifsud B, Tavares-Cadete F, Sugar R, Javierre B-M, Nagano T, Katsman Y, Sakthidevi M, Wingett SW, et al. 2015. The pluripotent regulatory circuitry connecting promoters to their long-range interacting elements. Genome Res. 25(4):582–597. 10.1101/gr.185272.114

Sessa A, Ciabatti E, Drechsel D, Massimino L, Colasante G, Giannelli S, Satoh T, Akira S, Guillemot F, Broccoli V. 2017. The Tbr2 Molecular Network Controls Cortical Neuronal Differentiation Through Complementary Genetic and Epigenetic Pathways. Cereb Cortex. 27(6):3378–3396. 10.1093/cercor/bhw270

Shen L, Shao N, Liu X, Nestler E. 2014. ngs. plot: Quick mining and visualization of next-generation sequencing data by integrating genomic databases. BMC Genomics. 15(1):1–14.

Shen Q, Wang Y, Dimos JT, Fasano CA, Phoenix TN, Lemischka IR, Ivanova NB, Stifani S, Morrisey EE, Temple S. 2006. The timing of cortical neurogenesis is encoded within lineages of individual progenitor cells. Nat Neurosci. 9(6):743–751. 10.1038/nn1694

Soleimani VD, Yin H, Jahani-Asl A, Ming H, Kockx CE, van Ijcken WF, Grosveld F, Rudnicki MA. 2012. Snail regulates MyoD binding-site occupancy to direct enhancer switching and differentiation-specific transcription in myogenesis. Mol Cell. 47(3):457–468.

Stark R, Brown G. 2011. DiffBind: differential binding analysis of ChIP-Seq peak data. R Package Version. 100(4.3).

Suh H, Consiglio A, Ray J, Sawai T, D’Amour KA, Gage FH. 2007. In Vivo Fate Analysis Reveals the Multipotent and Self-Renewal Capacities of Sox2+ Neural Stem Cells in the Adult Hippocampus. Cell Stem Cell. 1(5):515–528. 10.1016/j.stem.2007.09.002

Sun J, Rockowitz S, Xie Q, Ashery-Padan R, Zheng D, Cvekl A. 2015. Identification of in vivo DNA-binding mechanisms of Pax6 and reconstruction of Pax6-dependent gene regulatory networks during forebrain and lens development. Nucleic Acids Res. 43(14):6827–6846. 10.1093/nar/gkv589

Suter DM, Tirefort D, Julien S, Krause K-H. 2009. A Sox1 to Pax6 switch drives neuroectoderm to radial glia progression during differentiation of mouse embryonic stem cells. Stem Cells. 27(1):49–58. 10.1634/stemcells.2008-0319

Takemoto T, Uchikawa M, Kamachi Y, Kondoh H. 2006. Convergence of Wnt and FGF signals in the genesis of posterior neural plate through activation of the Sox2 enhancer N-1. Development. 133(2):297–306. 10.1242/dev.02196

Takemoto T, Uchikawa M, Yoshida M, Bell DM, Lovell-Badge R, Papaioannou VE, Kondoh H. 2011. Tbx6-dependent Sox2 regulation determines neural or mesodermal fate in axial stem cells. Nature. 470(7334):394–398. 10.1038/nature09729

Tang Z, Luo OJ, Li X, Zheng M, Zhu JJ, Szalaj P, Trzaskoma P, Magalska A, Wlodarczyk J, Ruszczycki B, et al. 2015. CTCF-Mediated Human 3D Genome Architecture Reveals Chromatin Topology for Transcription. Cell. 163(7):1611–1627. 10.1016/j.cell.2015.11.024

Taranova OV, Magness ST, Fagan BM, Wu Y, Surzenko N, Hutton SR, Pevny LH. 2006. SOX2 is a dose-dependent regulator of retinal neural progenitor competence. Genes Dev. 20(9):1187–1202. 10.1101/gad.1407906

Taylor T, Sikorska N, Shchuka VM, Chahar S, Ji C, Macpherson NN, Moorthy SD, Kort MAC de, Mullany S, Khader N, et al. 2022. Transcriptional regulation and chromatin architecture maintenance are decoupled functions at the Sox2 locus. Genes Dev. 36(11–12):699–717. 10.1101/gad.349489.122

Thompson JJ, Lee DJ, Mitra A, Frail S, Dale RK, Rocha PP. 2022. Extensive co-binding and rapid redistribution of NANOG and GATA6 during emergence of divergent lineages. Nat Commun. 13(1):4257.

Thomson M, Liu SJ, Zou L-N, Smith Z, Meissner A, Ramanathan S. 2011. Pluripotency factors in embryonic stem cells regulate differentiation into germ layers. Cell. 145(6):875–889.

Thurman RE, Rynes E, Humbert R, Vierstra J, Maurano MT, Haugen E, Sheffield NC, Stergachis AB, Wang H, Vernot B. 2012. The accessible chromatin landscape of the human genome. Nature. 489(7414):75–82.

Tobias IC, Abatti LE, Moorthy SD, Mullany S, Taylor T, Khader N, Filice MA, Mitchell JA. 2021. Transcriptional enhancers: from prediction to functional assessment on a genome-wide scale. Genome. 64(4):426–448. 10.1139/gen-2020-0104

Tolhuis B, Palstra R-J, Splinter E, Grosveld F, de Laat W. 2002. Looping and Interaction between Hypersensitive Sites in the Active β-globin Locus. Mol Cell. 10(6):1453–1465. 10.1016/S1097-2765(02)00781-5

Tomioka M, Nishimoto M, Miyagi S, Katayanagi T, Fukui N, Niwa H, Muramatsu M, Okuda A. 2002. Identification of Sox-2 regulatory region which is under the control of Oct-3/4–Sox-2 complex. Nucleic Acids Res. 30(14):3202–3213. 10.1093/nar/gkf435

Tuan DY, Solomon WB, London IM, Lee DP. 1989. An erythroid-specific, developmental-stage-independent enhancer far upstream of the human “beta-like globin” genes. Proc Natl Acad Sci. 86(8):2554–2558. 10.1073/pnas.86.8.2554

Uchikawa M, Ishida Y, Takemoto T, Kamachi Y, Kondoh H. 2003. Functional Analysis of Chicken Sox2 Enhancers Highlights an Array of Diverse Regulatory Elements that Are Conserved in Mammals. Dev Cell. 4(4):509–519. 10.1016/S1534-5807(03)00088-1

Uchikawa M, Kondoh H. 2016. Regulation of Sox2 via Many Enhancers of Distinct Specificities. In: Sox2. Elsevier; p. 107–129. 10.1016/B978-0-12-800352-7.00007-4

Villar D, Berthelot C, Aldridge S, Rayner TF, Lukk M, Pignatelli M, Park TJ, Deaville R, Erichsen JT, Jasinska AJ. 2015. Enhancer evolution across 20 mammalian species. Cell. 160(3):554–566.

Wamstad JA, Alexander JM, Truty RM, Shrikumar A, Li F, Eilertson KE, Ding H, Wylie JN, Pico AR, Capra JA, et al. 2012. Dynamic and Coordinated Epigenetic Regulation of Developmental Transitions in the Cardiac Lineage. Cell. 151(1):206–220. 10.1016/j.cell.2012.07.035

Webb AE, Pollina EA, Vierbuchen T, Urbán N, Ucar D, Leeman DS, Martynoga B, Sewak M, Rando TA, Guillemot F, et al. 2013. FOXO3 Shares Common Targets with ASCL1 Genome-wide and Inhibits ASCL1-Dependent Neurogenesis. Cell Rep. 4(3):477–491. 10.1016/j.celrep.2013.06.035

Weintraub AS, Li CH, Zamudio AV, Sigova AA, Hannett NM, Day DS, Abraham BJ, Cohen MA, Nabet B, Buckley DL, et al. 2017. YY1 Is a Structural Regulator of Enhancer-Promoter Loops. Cell. 171(7):1573–1588.e28. 10.1016/j.cell.2017.11.008

Welch RP, Lee C, Imbriano PM, Patil S, Weymouth TE, Smith RA, Scott LJ, Sartor MA. 2014. ChIP-Enrich: gene set enrichment testing for ChIP-seq data. Nucleic Acids Res. 42(13):e105–e105.

Wickham H, Averick M, Bryan J, Chang W, McGowan LD, François R, Grolemund G, Hayes A, Henry L, Hester J. 2019. Welcome to the Tidyverse. J Open Source Softw. 4(43):1686.

Xu X, Stoyanova EI, Lemiesz AE, Xing J, Mash DC, Heintz N. 2018. Species and cell-type properties of classically defined human and rodent neurons and glia. eLife. 7:e37551. 10.7554/eLife.37551

Ying Q-L, Stavridis M, Griffiths D, Li M, Smith A. 2003. Conversion of embryonic stem cells into neuroectodermal precursors in adherent monoculture. Nat Biotechnol. 21(2):183–186. 10.1038/nbt780

Yu G, Wang L-G, Han Y, He Q-Y. 2012. clusterProfiler: an R package for comparing biological themes among gene clusters. Omics J Integr Biol. 16(5):284–287.

Zappone MV, Galli R, Catena R, Meani N, Biasi SD, Mattei E, Tiveron C, Vescovi AL, Lovell-Badge R, Ottolenghi S, Nicolis SK. 2000. Sox2 regulatory sequences direct expression of a β-geo transgene to telencephalic neural stem cells and precursors of the mouse embryo, revealing regionalization of gene expression in CNS stem cells. Development. 127(11):2367–2382. 10.1242/dev.127.11.2367

Zeisel A, Hochgerner H, Lönnerberg P, Johnsson A, Memic F, van der Zwan J, Häring M, Braun E, Borm LE, La Manno G, et al. 2018. Molecular Architecture of the Mouse Nervous System. Cell. 174(4):999–1014.e22. 10.1016/j.cell.2018.06.021

Zhang J, Zhou Y, Yue W, Zhu Z, Wu X, Yu S, Shen Q, Pan Q, Xu W, Zhang R. 2022. Super-enhancers conserved within placental mammals maintain stem cell pluripotency. Proc Natl Acad Sci. 119(40):e2204716119.

Zhang Y, Liu T, Meyer CA, Eeckhoute J, Johnson DS, Bernstein BE, Nusbaum C, Myers RM, Brown M, Li W, Liu XS. 2008. Model-based Analysis of ChIP-Seq (MACS). Genome Biol. 9(9):R137. 10.1186/gb-2008-9-9-r137

Zhang Y, Wong C-H, Birnbaum RY, Li G, Favaro R, Ngan CY, Lim J, Tai E, Poh HM, Wong E, et al. 2013. Chromatin connectivity maps reveal dynamic promoter–enhancer long-range associations. Nature. 504(7479):306–310. 10.1038/nature12716

Zhou HY, Katsman Y, Dhaliwal NK, Davidson S, Macpherson NN, Sakthidevi M, Collura F, Mitchell JA. 2014. A Sox2 distal enhancer cluster regulates embryonic stem cell differentiation potential. Genes Dev. 28(24):2699–2711. 10.1101/gad.248526.114

Zhu LJ, Gazin C, Lawson ND, Pagès H, Lin SM, Lapointe DS, Green MR. 2010. ChIPpeakAnno: a Bioconductor package to annotate ChIP-seq and ChIP-chip data. BMC Bioinformatics. 11:1–10.

Ziller MJ, Edri R, Yaffe Y, Donaghey J, Pop R, Mallard W, Issner R, Gifford CA, Goren A, Xing J. 2015. Dissecting neural differentiation regulatory networks through epigenetic footprinting. Nature. 518(7539):355–359.

