## Supplemental data for "A *Sox2* Enhancer Cluster Regulates Region-Specific Neural Fates from Mouse Embryonic Stem Cells"

Short Title: Sox2 enhancers control neural identity

Ian C Tobias^1,2&^, Sakthi D Moorthy^1,3&^, Virlana M Shchuka^1^, Lida Langroudi^1^, Mariia Cherednychenko^1^, Zoe E Gillespie^1^, Andrew G Duncan^1^, Ruxiao Tian^1^, Natalia A Gajewska ^1^, Raphaël B Di Roberto^1^, and Jennifer A Mitchell^1*^

1) Department of Cell and Systems Biology, University of Toronto, Toronto, Ontario, M5S3G5, Canada

2) Current Address: Department of Biomedical Sciences, University of Guelph, Guelph, Ontario, N1G 2W1, Canada

3) Current Address: Center for Commercialization of Regenerative Medicine, Toronto, Ontario, M5G1M1, Canada

*Corresponding Author

 (J.A.M.)

^&^These authors made equal contributions to this work

**Supplemental Figures**

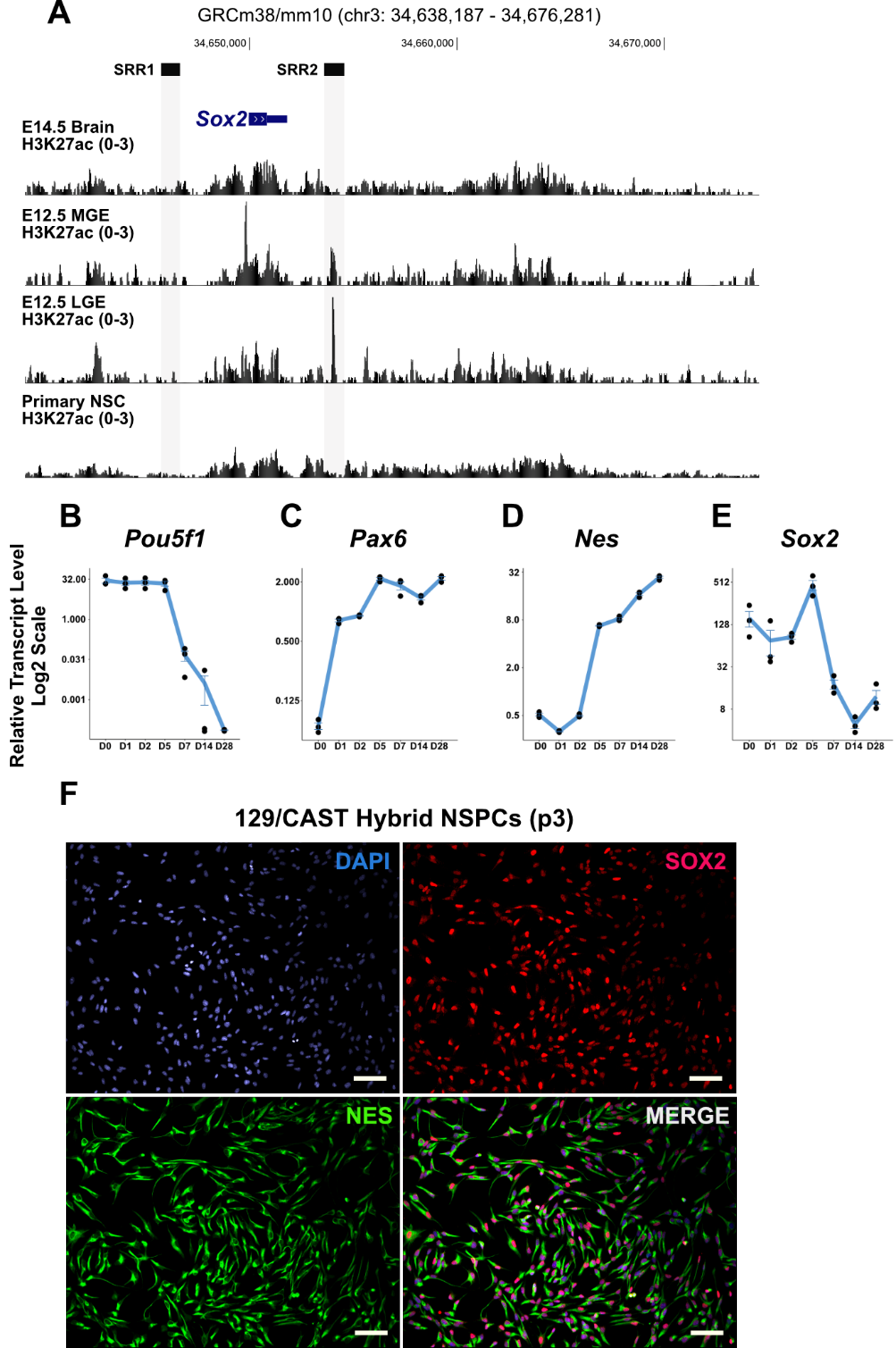

**Supplemental Figure S1. *Sox2* flanking region H3K27ac ChIP-seq profiles and neural lineage marker RT-qPCR.** A) H3K27ac ChIP-seq tracks visualized by UCSC Genome Browser (mm10). Shaded bars mark known Sox2 regulatory regions (SRRs) active in mouse embryonic stem cells. Regions of H3K27ac enrichment located further upstream (-8 kb) and downstream of *Sox2* (+10 to +16 kb) are enhancer candidates in the whole brain (E14.5), the E12.5 medial (MGE) and lateral (LGE) ganglionic eminences in cortical neurogenesis, and in isolated neural stem/progenitor cells (NSPCs). B) Pou5f1, C) Pax6, D) Nes, and E) Sox2 expression in differentiating F1 hybrid ESCs by RT-qPCR. Expression levels are shown relative to internal control transcripts Gapdh and Eef2a. Error bars represent standard deviation. n ≥ 3. (F) Immunofluorescence of microscopy of NES overlayed with SOX2 in F1 NSPCs. Scale bar distance is 50 µm.

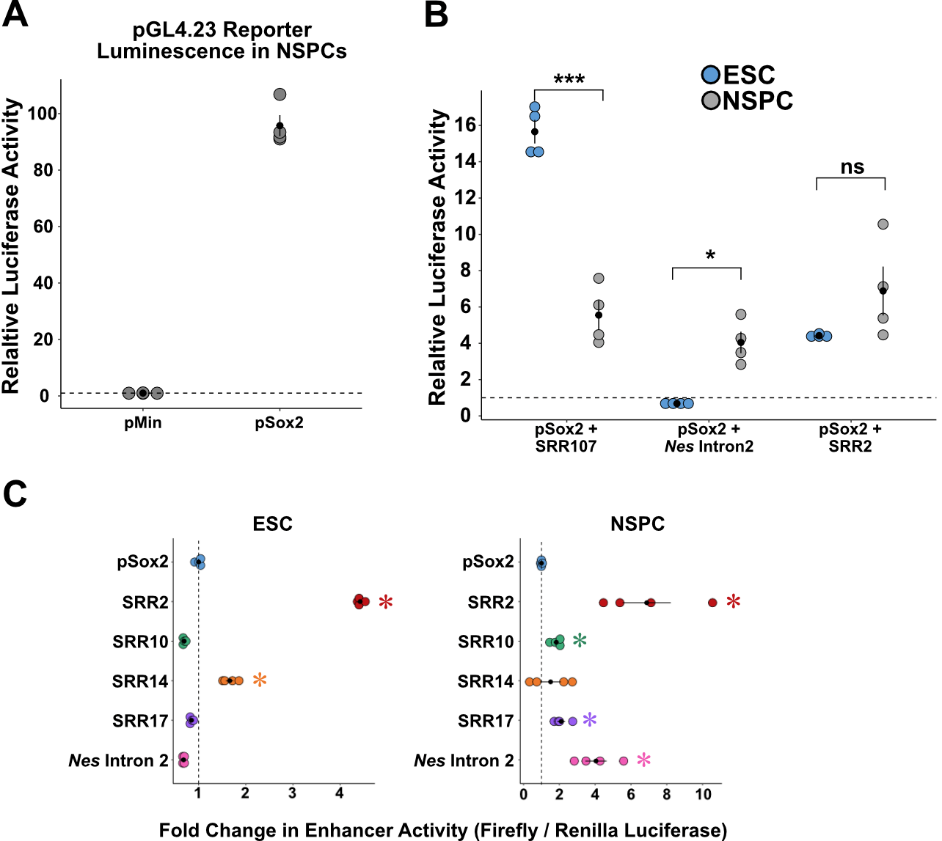

**Supplemental Figure S2. Dual luciferase reporter assay analysis of undifferentiated ESCs versus ESC-generated NSPCs.** A) Luciferase reporter activity values following F1 NSPC transfection with the original minimal promoter (pMin) or modified with the Sox2 plasmid by restriction cloning. B) Fold increase in luciferase reporter activity in F1 ESCs and F1 NSPCs with enhancer insertion at the NotI restriction site. Dashed line represents empty pSox2 vector luciferase activity values. Enhancers are Sox2 regulatory region 107 (SRR107), Nestin intron 2 enhancer, and SRR2. Error bars represent SEM. n ≥ 3. (∗) P < 0.05, (∗∗) P < 0.01, (∗∗∗) P < 0.001, (∗∗∗∗) P < 0.0001, (ns) not significant.

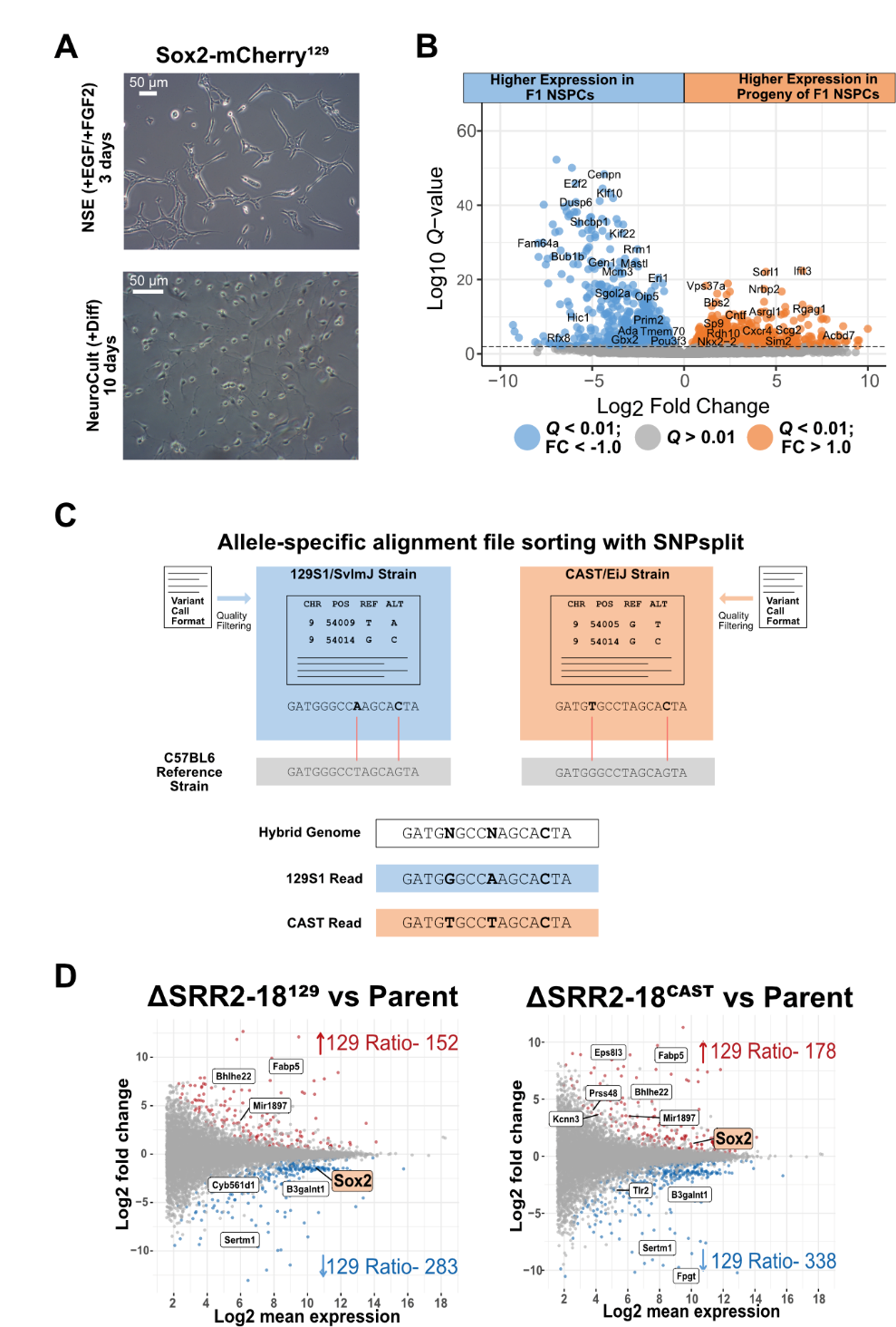

**Supplemental Figure S3. Representative morphology and gating of Sox2-mCherry positive NSPCs with genome-wide allele-specific RNA-seq read sorting approach.** A) Representative morphologies in cultured cells by phase contrast microscopy. F1 hybrid embryonic stem (ES) and neural stem/progenitor (NPS) cells with a M. musculus¹²⁹ (129) x M. Castaneous (Cast) background are shown in maintenance culture conditions. B) Volcano plot and the number of differentially expressed genes between the F1 NSPCs and the progeny of the same cells after 10 days of undirected differentiation. The colored data points show the transcripts passed the cutoffs of absolute log2 fold-change > 1 and Q < 0.01. C) Schematic of computational approach to sort next-generation sequencing alignment data overlapping single nucleotide polymorphisms between the 129S and Cast sub-strains N-masked in the custom hybrid genome assembly. D) MA plots showing significant allelic imbalances in allele-sorted RNA-seq data comparing ΔSRR2-18^129^ versus parent NSPCs and ΔSRR2-18^Cast^ versus parent NSPCs. Known and predicted genes on chromosome 3 are labeled.

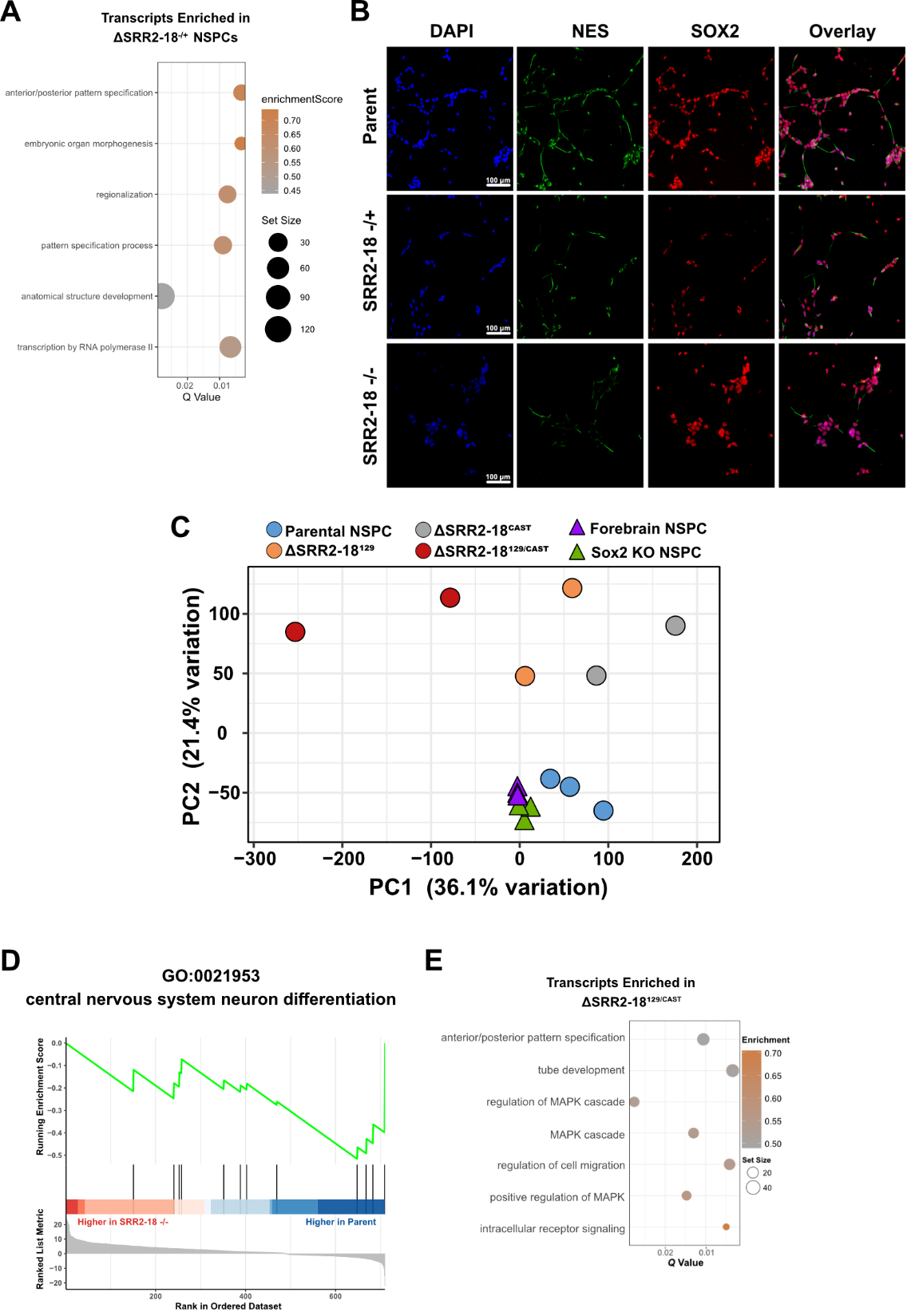

**Supplemental Figure S4. Gene set enrichment analyses in SRR2-18 deleted NSPCs with differential expression reanalysis of *Sox2* knockout neurospheres.** A) Bubble plot of the enriched GO biological process terms of genes nearest to the chromatin regions showing increased accessibility in SRR2-18^129/Cast^ vs. control NSPCs. B) Qualitative SOX2 and NES immunofluorescence micrographs of parent NSPCs, ΔSRR2-18 heterozygous, and ΔSRR2-18 homozygous NSPCs. Scale bar is 100 µm. C) Biplot of principle component analysis comparing RNA-seq data of parent and enhancer-deleted NSPCs to wild-type and Sox2 knockout (KO) neurosphere RNA-seq data compiled from the GEO repository D) Enrichment plot showing depletion of genes annotated by gene ontology identifier GO:0021953 (central nervous system neuron differentiation) in SRR2-18^129/Cast^ vs. control NSPCs. E) Bubble plot of the significantly enriched gene sets of upregulated genes in ∆SRR2-18^129/Cast^ vs. control NSPCs.

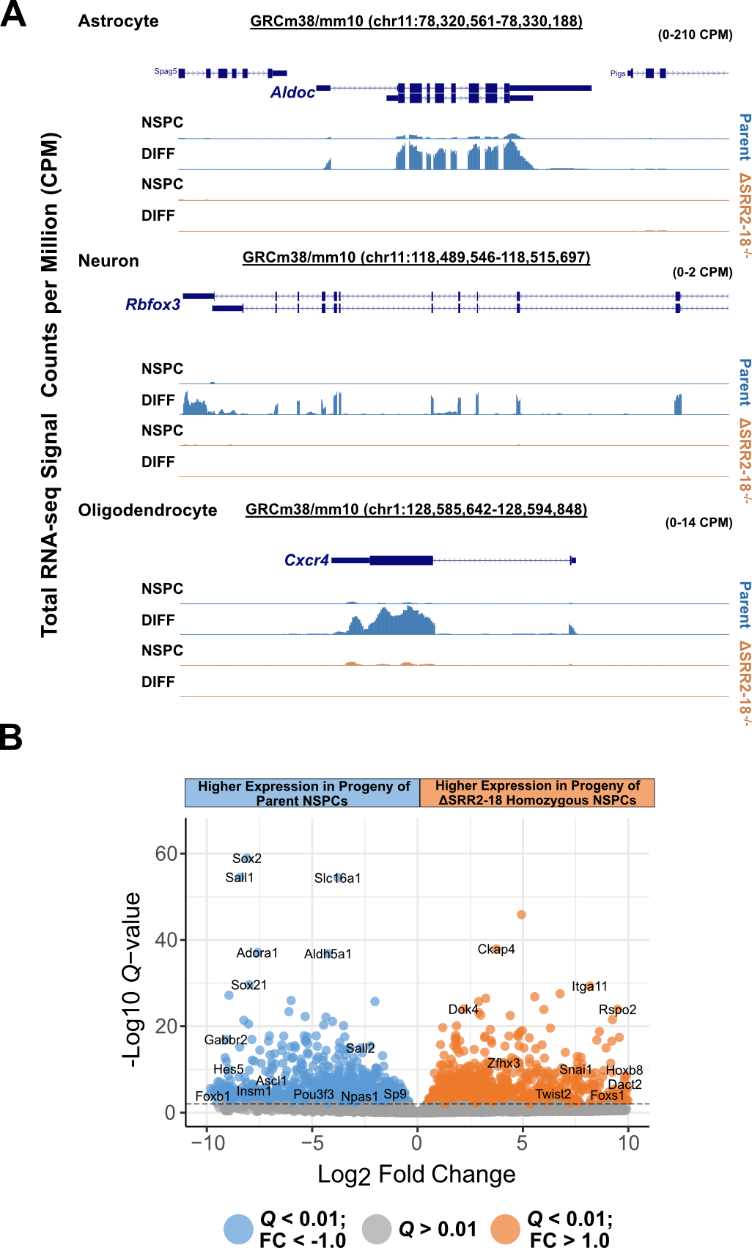

**Supplemental Figure S5. Differential expression analysis of differentiated progenies of SRR2-18 homozygous deletion versus parental NSPCs.** A) Normalized RNA-seq tracks over the *Aldoc* (astrocyte marker gene), *Rbfox3/NeuN* (neuronal marker gene), and *Cxcr4* (oligodendrocyte marker gene) loci, displayed on the UCSC Genome Browser (mm10). Signal tracks show greater transcript read coverage in libraries produced from the differentiated progeny of parent NSPCs to that of SRR2-18^129/Cast^ NSPC derivatives. B) Volcano plot and the number of differentially expressed genes between the differentiated progeny of parent versus ΔSRR2-18^129/Cast^ cells. Colored data points show the transcripts passed the cutoffs of absolute log2 fold-change > 0.5 and Q < 0.01. Q values represent the adjusted P values computed with the Benjamini & Hochberg method for controlling false discovery rate (FDR).

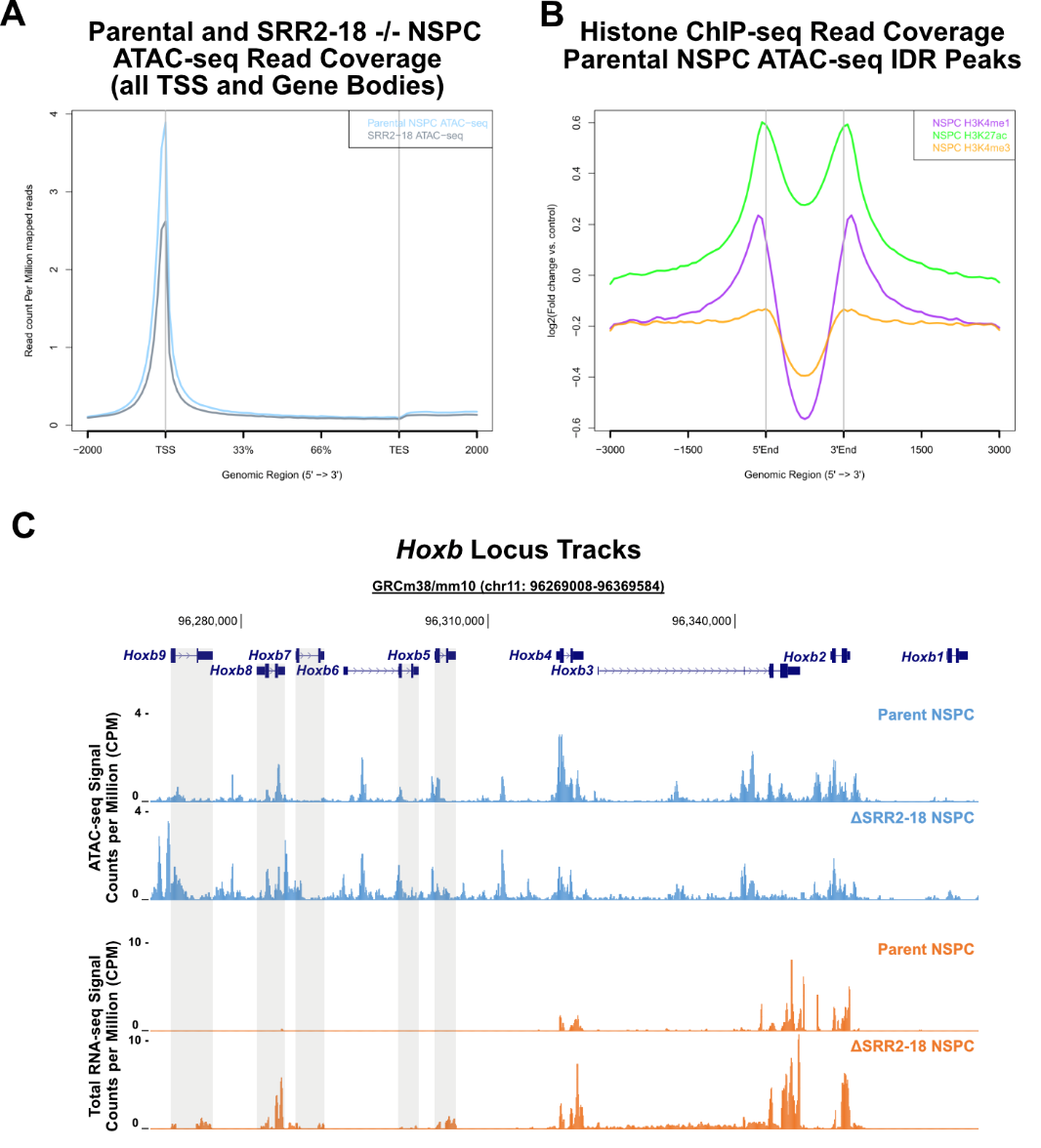

**Supplemental Figure S6. NSPC ATAC-seq read coverage over mouse genes, enhancer chromatin features, and Hoxb locus accessible chromatin profile.** A) Normalized ATAC-seq read coverage at the transcription start site (TSS), gene body and flanking regions in parent and SRR2-18^-/-^ NSPCs. B) Normalized ChIP-seq read coverage for histone post-translational modifications (H3K27ac, H3K4me1, H3K4me3) associated with regulatory DNA across parent NSPC ATAC-seq peaks. C) Normalized ATAC-seq and RNA-seq tracks over the *Hoxb* locus, displayed on the UCSC Genome Browser (mm10). Signal tracks show increased accessibility at the *Hoxb6-9* genes and increased total *Hoxb6-9* RNA in ΔSRR2-18^129/Cast^ NSPCs.

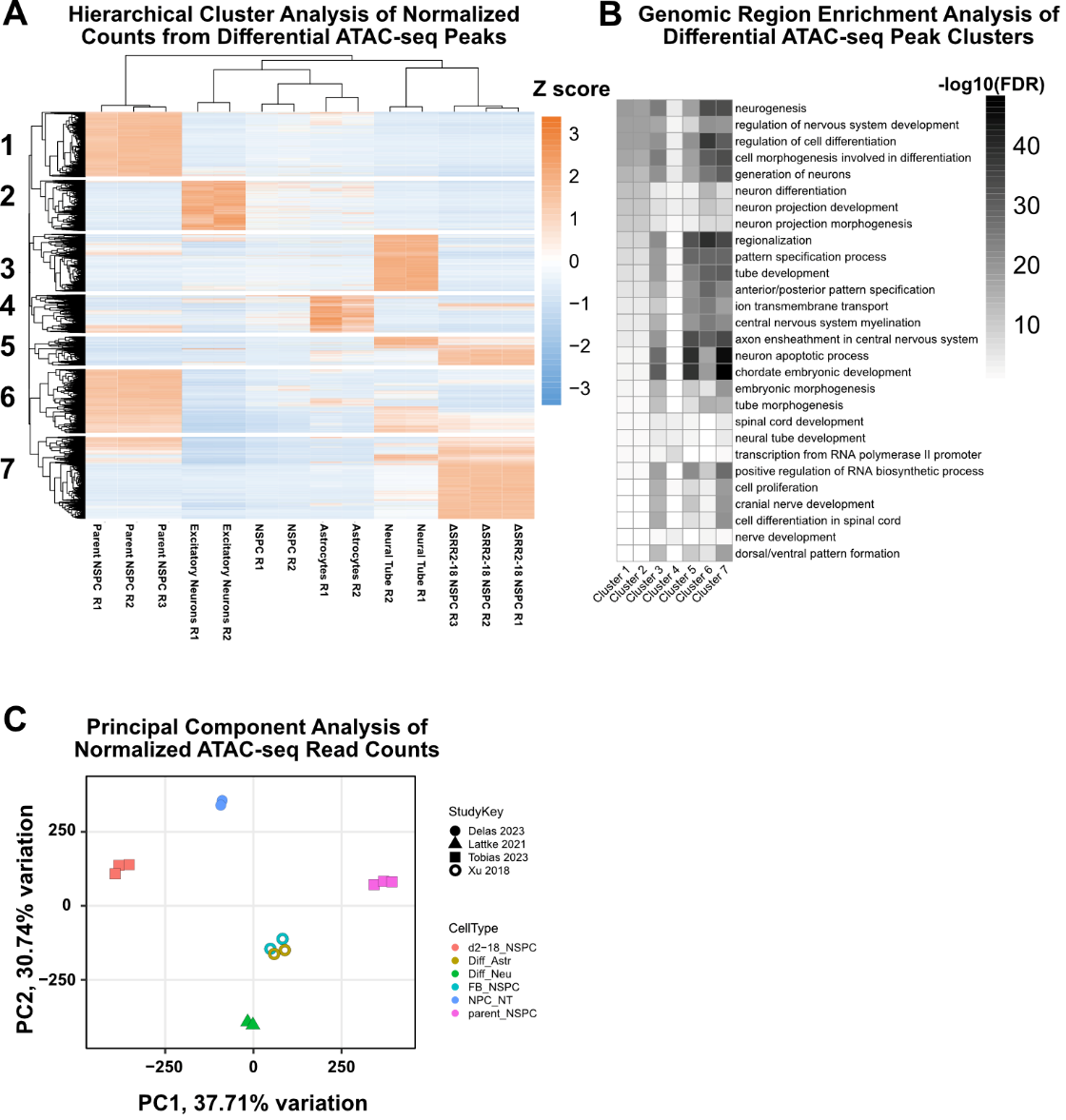

**Supplemental Figure S7. Accessible genomic region hierarchical cluster analysis and cluster annotation by genomic region enrichment analysis.** A) Hierarchical cluster analysis of normalized counts from differential ATAC-seq peaks. Clustering revealed groups of genomic regions with differential accessibility that distinguished Sox2-tagged F1 NSPCs (parent) from ΔSRR2-18^129/Cast^ , forebrain derived neural progenitors (NSPC), astrocyte (ASTR), excitatory neurons (NEU), and neural tube progenitors (NPC NT). B) GO biological process term enrichment across differentially accessible regions that define clusters 1-7. C) Principal component analysis of normalized ATAC-seq read counts showing that most datasets analyzed are well separated aside from NSPCs and astrocyte accessible chromatin signatures.

**Supplemental Tables**

**Table S1.** Raw sequencing data compiled from the ENCODE consortium.

**Table S2.** Raw sequencing data compiled from NCBI Sequence Read Archive (SRA) and European Nucleotide Archive (ENA)

**Table S3.** Proximal Sox2 regulatory region coordinates.

**Table S4.** Differential gene expression analysis of in vitro differentiated parent NSPCs versus undifferentiated control.

**Table S5.** Differential gene expression analysis of SRR2-18 heterozygous deletion versus control.

**Table S6.** Gene set enrichment analysis for GO biological process terms in differentially abundant transcripts in SRR2-18 heterozygous deletion versus control.

**Table S7.** Differential gene expression analysis of SRR2-18 homozygous deletion versus control.

**Table S8.** Gene set enrichment analysis for GO biological process terms in differentially abundant transcripts in SRR2-18 homozygous deletion versus control.

**Table S9.** Differential gene expression analysis of differentiated progenies from SRR2-18 NSPCs homozygous deletion versus control.

**Table S10.** Gene set enrichment analysis for GO biological process terms in differentially abundant transcripts in differentiated progenies from SRR2-18 NSPCs homozygous deletion versus control.

**Table S11.** Differential chromatin accessibility analysis of SRR2-18 NSPCs homozygous deletion versus control.

**Table S12.** List of non-redundant SOX2 bound regions in neural progenitors from reanalyzed SOX2 ChIP-seq data.

**Table S13.** JASPAR motif matches recovered from intersectional SOX2 ChIP-seq, H3K27ac-modified and accessible chromatin regions in NSPCs.

**Table S14.** Genomic region enrichment analysis for GO biological process terms depleted in differentially accessible regions in SRR2-18 NSPCs homozygous deletion versus control.

**Table S15.** Genomic region enrichment analysis for GO biological process terms enriched in differentially accessible regions in SRR2-18 NSPCs homozygous deletion versus control.

**Table S16.** List of guide RNA (gRNA) target sequences

**Table S17.** List of primers for plasmid construction

**Table S18.** List of antibodies

**Table S19.** List of RT-qPCR primers
